# Snf1p/Hxk2p/Mig1p pathway regulates exponential growth, mitochondrial respiration, and hexose transporters transcription in *Saccharomyces cerevisiae*

**DOI:** 10.1101/2020.06.22.165753

**Authors:** Andres Carrillo-Garmendia, Cecilia Martinez-Ortiz, Jairo Getzemani Martinez-Garfias, Juan Carlos González-Hernández, Gerardo M. Nava, Minerva Ramos-Gomez, Miguel David Dufoo-Hurtado, Luis Alberto Madrigal-Perez

## Abstract

The Crabtree effect occurs under high-glucose concentrations and is characterized by the increase of the growth and a decrease in mitochondrial respiration of yeasts. Regulation of the Crabtree effect could enhance ethanol production with biotechnological purposes and a better understanding of the etiology of cancer due to its similitude with the Warburg effect. Nonetheless, the conclusive molecular mechanism of the Crabtree effect is still on debate. The pathway Snf1p/Hxk2p/Mig1p has been linked with the transcriptional regulation of the hexose transporters and has also been identified in the modulation of phenotypes related to the Crabtree effect. Nevertheless, it has not been directly identified the genetic regulation of the hexose transporters with modulation of the Crabtree effect phenotypes by Snf1p/Hxk2p/Mig1p pathway. In this sense, we provide evidence that the deletion of the *SNF1* and *HXK2* genes affects the exponential growth, mitochondrial respiration, and the transcription of hexose transporters in a glucose-dependent manner in *Saccharomyces cerevisiae*. The *Vmax* of the main hexose transporters transcribed showed a positive correlation with the exponential growth and a negative correlation with the mitochondrial respiration. Transcription of the gene *HXT2* was the most affected by the deletion of the pathway *SNF1/HXK2/MIG1*. Deletion of the orthologous genes *SNF1* and *HXK2* in the Crabtree negative yeast, *K. marxianus,* has a differential effect in exponential growth and mitochondrial respiration in comparison with *S. cerevisiae*. Overall, these results indicate that the *SNF1/HXK2/MIG1* pathway transcriptionally regulates the hexose transporters having an influence in the exponential growth and mitochondrial respiration in a glucose-dependent manner.

## Introduction

The metabolic flexibility of organisms to form ATP under different environmental circumstances is a valuable phenotype to perpetuating its species. In the case of *Saccharomyces cerevisiae*, it has evolved an intricate signaling pathway to adapt its metabolism depending on the carbohydrate accessibility to form ATP by fermentation (substrate-level phosphorylation) or by mitochondrial respiration (oxidative phosphorylation) [1, 2]. Sugar concentration above 0.8 mM activates a metabolic pathway called glucose repression that reshapes the metabolism to produce ATP mainly by fermentation; accompanied by an increase in the glycolytic flux, decreasing the mitochondrial respiration and increasing the fermentation metabolites, altogether this phenotype is called Crabtree effect. However, the molecular basis behind the Crabtree effect is still unknown. Some hypotheses point out that the glycolysis pathway is the key step during the Crabtree effect. For example, the increase of the glycolysis-derived hexose phosphate, fructose-1,6-bisphosphate inhibits the respiratory chain in isolated mitochondria of *S. cerevisiae* [3]. Additionally, glucose supplementation in AS-30D hepatoma cells decreases intracellular phosphate (P_i_) and increases the AMP, glucose-6-phosphate, fructose-6-phosphate, and fructose-1,6-bisphosphate [4]. Interestingly, when isolated mitochondria were incubated with low P_i_ (0.6 mM), inhibition of the mitochondrial respiration was also observed [4]. Besides, it has been shown that the augmenting of glucose concentration in *S. cerevisiae* increases the glycolytic flux accompanied by inhibition of mitochondrial respiration [5]. Altogether, these data indicate that an increase in the glycolytic pathway flux is important to unleash the Crabtree effect. Nonetheless, which signal/pathway regulates this increase in glycolytic flux is still unknown. One hypothesis suggests that hexose transporters could regulate the glycolytic flux. For instance, in a *S. cerevisiae* strain in which all the hexose transporters were deleted, the expression of a low-affinity transporter Hxt1p (*Km* around 90 mM and *Vmax* ~ 690 nmol min^−1^ mg^−1^ [6]) have a major glycolytic flux than the expression of the high-affinity transporter Hxt7p (*Km* ~ 1 mM and *Vmax* ∼ 186 nmol min^−1^ mg^−1^ [6]) [7]. Interestingly, the diminution in the glycolytic flux resulted in a decrease in the ethanol accumulation accompanied by an increase in the mitochondrial respiration [7]. It has been proposed that the glucose uptake has a linear correlation with the ethanol production in *S. cerevisiae*, which implies that hexose transporters could play an important role in the switch between fermentation and mitochondrial respiration [8]. The hexose transporters have different biochemical properties. Thereby, the expression of the hexoses transporter occurs under different sugar concentrations and fits with each environmental situation. Nevertheless, the signaling pathway that modulates those changes in the hexose transporters expression is not well known.

The protein Snf1p is AMP-activated protein kinase (orthologous to the mammalian AMPK) that responds to energetic changes mainly through the ADP/ATP ratio in *S. cerevisiae* [9]. Interestingly, the deletion of the gene *SNF1* causes a severe metabolic disruption in the response to glucose concentrations [10–12]. For example, *SNF1* deletion reverses the mitochondrial respiration inhibition in high-glucose concentrations [10]. Snf1p regulates by phosphorylation the translocation of the nucleus of two important regulators of glucose repression: Hxk2p [13] and Mig1p [14]. Hxk2p has a dual function, as catalytic in the first reaction of glycolysis pathway and as a transcriptional regulator. Hx2p also serves as an intracellular glucose sensor changing its conformation to interact with Mig1p under high-glucose concentrations to repress *SUC2* promoter [15]. Even, Hxk2p is necessary for the inhibition of the Mig1p by Snf1p, when both form part of the repressor complex [13]. Mig1p is a Cys_2_-His_2_ zinc finger protein that binds to glucose-repressible genes and its localization in the nucleus is dependent on the glucose concentration. Interestingly, Mig1p regulates transcriptionally the promoters of the genes *HXT2* and *HXT4* preventing its transcription at 4% glucose [16]. Indeed, the deletion of the gene *MIG1* increases the transcription of the hexose transporters *HXT3*, *HXT4*, *HXT6*, and *HXT7* at 4% glucose; while the transcription of the gene *HXT8* was decreased at the same conditions [17]. Besides, transcription of the gene *HXT4* was augmented in the *mig1*Δ strain at 2% and 10% glucose [18].

For this reason, the aim of this study was to elucidate if the pathway Snf1p/Mig1p/Hxk2p is responsible for the regulation of transcription of hexose transporters in a glucose-dependent manner and its relationship with exponential growth and mitochondrial respiration in yeasts. The data provided in this paper indicate that *S. cerevisiae* regulates the transcription of the hexose transporters in a glucose-dependent manner by the pathway Snf1p/Mig1p/Hxk2p. Specifically, the transcription of the high-affinity *HXT2* was several affected by the deletion of the genes *SNF1*, *HXK2*, and *MIG1*. Importantly, we also found that exponential growth correlates positively with the *Vmax* of the mainly transcribed hexose transporters and correlates negatively with the basal respiration. These data indicate that hexose transporters might have a relevant impact on exponential growth and mitochondrial respiration and are transcriptionally regulated by the pathway Snf1p/Mig1p/Hxk2p.

## Material and methods

### *Saccharomyces cerevisiae* strains

For performing this study, it was used the *S. cerevisiae* strain BY4741 (*MATa*; *ura3Δ0; leu2Δ0; his3Δ1; met15Δ0*) and its mutants in the genes *SNF1* (*snf1*Δ, pHLUM; *MATa; ura3Δ0//URA3; leu2Δ0//LEU2; his3Δ0//HIS3; met15Δ0//MET15*; YDR477w∷kanMX4), *HXK2* (*hxk2*Δ, *MATa*; *ura3Δ0; leu2Δ0; his3Δ1; met15Δ0*; YGL253w∷kanMX4), and *MIG1* (*mig1*Δ, *MATa; ura3Δ0; leu2Δ0; his3Δ1; met15Δ0*; YGL035c∷kanMX4). Also some experiments were performed with the strain BY4742 (*MATα; his3Δ1; leu2Δ0; lys2Δ0; ura3Δ0*) and its mutant in the gene *SNF1* (*snf1*Δ, *MATα; his3Δ1; leu2Δ0; lys2Δ0; ura3Δ0*; YDR477w:kanMX4). The *S. cerevisiae* strains were maintained in YPD medium (1% yeast extract, 2% casein peptone, and supplemented with 2% glucose), deletant strains were supplemented with geneticin (G-418 disulfate salt solution, Sigma-Aldrich) at a final concentration of 200 μg/mL.

### *Kluyveromyces marxianus* deleted strains

*K. marxianus* L-2029 strain was used to obtain deletions in the genes *SNF1* and *RAG5* orthologous to the *S. cerevisiae* genes *SNF1* and *HXK2*, respectively. The genes *SNF1* and *RAG5* were interrupted by homologous recombination using a PCR-based gene deletion strategy [19]. First, the *kanMX4* (to confer resistance to geneticin) gene was amplified with the primers: forward 5’-ATGCGTACGCTGCAGGTCGAC-3’ and reverse 5’-TCAATCGATGAATTCGAGCTCG-3’ [20]. In the second instance, a 45mers primers flanking upstream and downstream the open reading frame (ORF) of the interest gene (*SNF1* and *RAG5*) and with homology to the *kanMX4* were used. *SNF1* primers: forward 5’-AAAAAGGGGAAACAACAGAGAGATACAACAATTAAATTGGACATG-3’ and reverse 5’-TTACGTAATGAACTAACCGGTATGGAAGAGAAACACAAAAACTCA-3’. *RAG5* primers: forward 5’-GGTTTACGCCTCCAATTATTACAAACACACACACTTACAAAAATG-3’ and reverse 5’-CTTCGATACCAACAGACTTACCGGCAGCCAATCTCTTTTGTGTCA-3’. The PCR products were inserted in *K. marxianus* using lithium acetate transformation protocol described by Gietz, St Jean, Woods and Schiestl [21] with some modifications. Briefly, *K. marxianus* was grown in a 250 mL shake-flask with 50 mL of YPD media at 30 °C and 250 rpm until it reaches the mid-log phase (O.D_.600._ ~0.6). Cells were harvest at 4000 x *g* for 2 minutes at room temperature and washed two times with distilled water. Afterwards, cells were washing with TE/LiAc (10x TE; 0.1 M Tris-HCl, 0.01 M EDTA, pH 7.5; 10x lithium acetate (LiAc); 1 M LiAc pH 7.5, adjusted with diluted acetic acid) and resuspended in 1 mL 1x TE/LiAc. Cells transformation was carried out in a 1.5 mL microtube with 50 μL cells, 12 μL PCR product, 50 μg salmon sperm DNA, and 300 μL PEG 4000 solution (40% de PEG 4000, 1x TE, 1x LiAc). Microtube was incubated at 30 °C for 30 minutes and 100 rpm, follow by a termic shock in a water bath at 42 °C for 15 minutes, cells were harvest at 13000 x *g* for 5 seconds and resuspended in 1 mL 1x TE solution. Finally, transformant cells were selected in a YPD plate with 200 μg/mL geneticin (G-418, Sigma-Aldrich).

### Over-expression of the gene *HXK2*

The pYES2.1 TOPO TA yeast expression kit (Invitrogen) was used to over-express the gene *HXK2*. Primers to amplify the ORF of the gene *HXK2* were designed to obtain the ORF without stop codon. *HXK2* primers: forward 5′-ATGGTTCATTTAGGTCCAAAAAAACC-3′ and reverse 5′-AGCACCGATGATACCAACGGAC-3′. PCR reactions contained: 1 μL primers at 10 mM, 1 μL dNTP mix at 10 mM, 5 μL Dream Taq 10x buffer, 0.25 μL Dream Taq, 100 ng of *S. cerevisiae* genomic DNA, and 40.75 μL nuclease-free water. Amplicons were purified using PureLink PCR purification kit (Invitrogen). The plasmid constructs were cloned into *Escherichia coli* TOP10 (F^−^ *mcr*A Δ(*mrr*-*hsd*RMS-*mcr*BC) φ80*lac*ZΔM15 Δ*lac*X74 *rec*A1 *ara*D139 Δ(*ara-leu*)7697 *gal*U *gal*K λ^−^ *rps*L(Str^R^) *end*A1 *nup*G) by chemical transformation, which served as a bacterial host for the recombinant plasmid. Finally, yeast strains were transformed using LiAc method [22].

### Growth kinetics

The growth kinetics were carried out as described by Olivares-Marin, Madrigal-Perez, Canizal-Garcia, Garcia-Almendarez, Gonzalez-Hernandez and Regalado-Gonzalez [5]. Assay of overexpression in *S. cerevisiae* come from pre-inoculum grown with SC media (6.7 g/L yeast nitrogen base without amino acids, 2 g/L yeast synthetic drop-out media supplements without uracil, 0.05 g/L uracil, glucose according to the experiment), while growth kinetics of *Kluyveromyces marxianus* and *S. cerevisiae* were made with pre-inoculums grown in YPD media. Growth kinetics were started at O.D._600_ •0.01 in a 96-well plate that contained 200 μL of YPD media supplemented with 0.005% or 5% glucose. The 96-well plate was incubated at 30 °C for 15 hours with constant shaking. The growth was monitored measuring the O.D._600_ each hour using a microplate reader (Multiskan Go; Thermo-Scientific). Data were analyzed with the statistical package GraphPad Prism 6.00 for Macintosh (GraphPad Software) fitting the growth kinetics with the exponential growth equation to obtain the specific growth rate (*μ*).

### Mitochondrial respiration analyses

The *in situ* mitochondrial respiration was performed as described by Tello-Padilla, Perez-Gonzalez, Canizal-Garcia, Gonzalez-Hernandez, Cortes-Rojo, Olivares-Marin and Madrigal-Perez [23]. The pre-inoculum cultures were made in SC medium supplemented with 2% glucose and were used to select the recombinant strains (without uracil) and deleted strains (with geneticin). Then, the strains were cultured in 250 mL shake-flasks with 50 mL of YPD medium supplemented with 0.005% or 5% glucose at 30 °C and 250 rpm until the mid-log phase (OD_600_~0.5) was reached. Next, cells were harvested at 4000 x *g* for 5 minutes and washed three times with deionized water, and resuspended in a 1:1 ratio (*w/v*). Immediately, cells were ready to measure the oxygen consumption polarographically with a Clark type electrode connected to a YSI5300A monitor (Yellow Springs, OH, USA) and a computer for data acquisition. In the polarograph chamber were added: 125 mg of cells (wet weight), 5 mL of MES-TEA buffer (10 mM morphoethanolsulfonic acid, pH 6.0 with triethanolamine) and glucose (10 mM); the cells were maintained with constant agitation (basal respiration). Afterward, 0.015 mM carbonylcyanide-*p*-trichlorophenylhydrazone (CCCP) was added to the chamber to stimulate the uncoupled state (maximum respiration) and oxygen consumption was recorded for 3 minutes. Finally, inhibitors of the electron transport chain (ETC) were added: thenoyltrifluoroacetone (TTFA, 1 mM), antimycin A (AA), and 0.75 mM KCN (non-mitochondrial respiration), each inhibitor was left for 3 minutes. The results were analyzed using the statistical package GraphPad Prism 6.00 for Macintosh (GraphPad Software).

### RT-qPCR

The strains (BY4741 genetic background) were grown in 25 mL shake-flask with 5 mL of SC medium supplemented with 0.005% or 5% glucose until the mid-log phase (OD_600_~0.5) was reached. The second experiment was made with the strains with a genetic background BY4742, which were grown in 25 mL shake-flask with 5 mL of YPD medium supplemented with 0.01% or 5% glucose until the mid-log phase (OD_600_~0.5) was reached. Cells were harvested at 13000 x *g* for 3 minutes. TRIzol reagent (Ambion, Life Technologies) was used to isolated the total RNA following the manufacturer’s instructions. The RNA pellet was resuspended in 50 μL diethylpyrocarbonate water (DEPC), adding recombinant DNAase I, RNAse free (Roche Diagnostics GmbH). The integrity of the RNA was verified with an electrophoretic analysis, as well as the concentration and quality of the RNA using a μDrop plate in the Multiskan Go spectrophotometer (Thermo-Scientific).

To carry out the qPCR the cDNA synthesis was performed using 2.5 μg of total RNA, 200 ng of random hexamer primers, and 0.65 mM of dNTPs. The reaction mix was incubated at 65 °C for 5 minutes. Then, to reaction were added 4 μL of 5X buffer of RevertAid and 2 μL of DTT 0.1 M and incubated at 37 °C for 2 minutes. Finally, 200 U of the RevertAid RT was added, followed by two periods of incubation, 10 minutes at room temperature and at 37 °C for 50 minutes. Each qPCR reaction had a final volume of 10 μL, 1 μL of cDNA, 5 μL of SYBR Select Master Mix (Applied Biosystems), 0.4 μL of each primer and 3.2 μL of DEPC water were added. Subsequently, the PCR was performed in a Rotor-Gene Q thermocycler (Qiagen). Absolute quantification was used to quantify the transcription using as a reference gene *UBC6* (ubiquitin-conjugation-6) (**Table 1**). Conditions of the PCR were: activation of UDG at 50 °C for 2 minutes, followed by the activation of AmpliTaq-DNA at 95 °C for 2 minutes with 35 cycles of denaturation at 94 °C for 15 seconds. The alignment temperature of primers *HXT1, HXT3, HXT4, HXT6, HXT7*, and *HXT10* was 50 °C, for *HXT2* was 62 °C, and for *UBC6* was 51 °C for 30 seconds, and the extension temperature of 72 °C for 1 minute, followed by 1 cycle at 72 °C for 7 minutes.

**Table 1.**
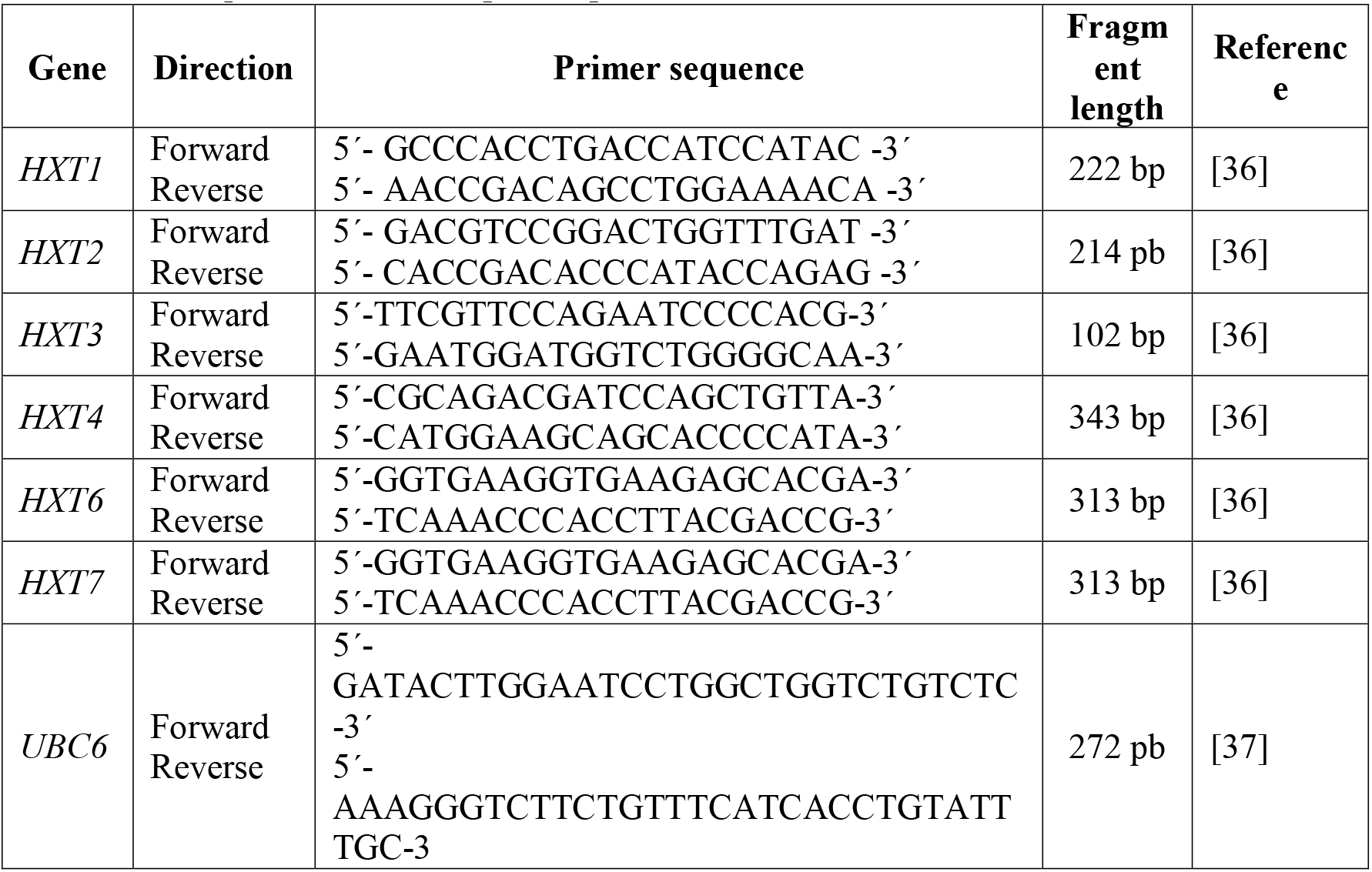
RT- qPCR hexose transporters primers.

### Statistical analyses

Data were obtained from at least three independent experiments and the mean ± standard deviation was graphed. Means were compared using one-way ANOVA followed by a Dunnett multiple comparisons to analyze differences in exponential growth and mitochondrial respiration. Differences between means in hexose transporters transcription were tested using a two-tailed unpaired *t*-test. Pearson’s correlation analysis was carried to evaluate the relationship between exponential growth/mitochondrial respiration and *Vmax*. Statistical analyses were computed in the software GraphPad Prism 6.00 for Macintosh (GraphPad Software).

Principal component analysis (PCA) plot was generated using ClustVis: (http://biit.cs.ut.ee/clustvis) utilizing a probabilistic PCA method without row scaling and employing the function (In x+1) for transformation [24].

## Results

### Effect of *HXK2* and *SNF1* deletion on the growth of *S. cerevisiae*

The transduction signal of the Snf1p/Hxk2p pathway responds primarily to glucose, impacting in the form that cells produce ATP. Exponential growth is tightly coupled to the form to obtain ATP in *S. cerevisiae*, showing higher rates of growth when having a fermentative growth in comparison when having respiratory growth [5]. We hypothesize that hexose transporters are the principals responsible for reshaping the *S. cerevisiae* cells to switch between mitochondrial respiration and fermentation, regulating the glycolytic flux. Taken those ideas into account, we propose that *S. cerevisiae* exponential growth has a correlation with the biochemical characteristics of the hexose transporters mainly transcribed, due to the mode to form ATP. For this reason, we decided to measure the exponential growth of the deleted strains *snf1*Δ and *hxk2*Δ under two concentrations of glucose: 0.005% (0.277 mM) and 5% (277 mM) that induce mitochondrial respiration and fermentation, respectively [5]. In this regard, a decrease in the specific growth rate is shown at 0.005% glucose (respiratory) in comparison to the growth at 5% glucose (fermentative) in the BY4741 strain (**Fig. 1a**), which corroborates the difference in the growth between a fermentative and respiratory glucose concentration. At 0.005% glucose, an increase in the specific growth rate occurs with the deletion of the genes *SNF1* and *HXK2* (**Fig. 1b**). However, only the strain *hxk2*Δ decreases the specific growth rate at 5% glucose (**Fig. 1c**). These results highlight the importance of *SNF1* in the growth at low-glucose concentrations that perform respiratory growth. Interestingly, *HXK2* gene deletion has a dual effect on growth dependently of the glucose concentration.

**Fig 1.**
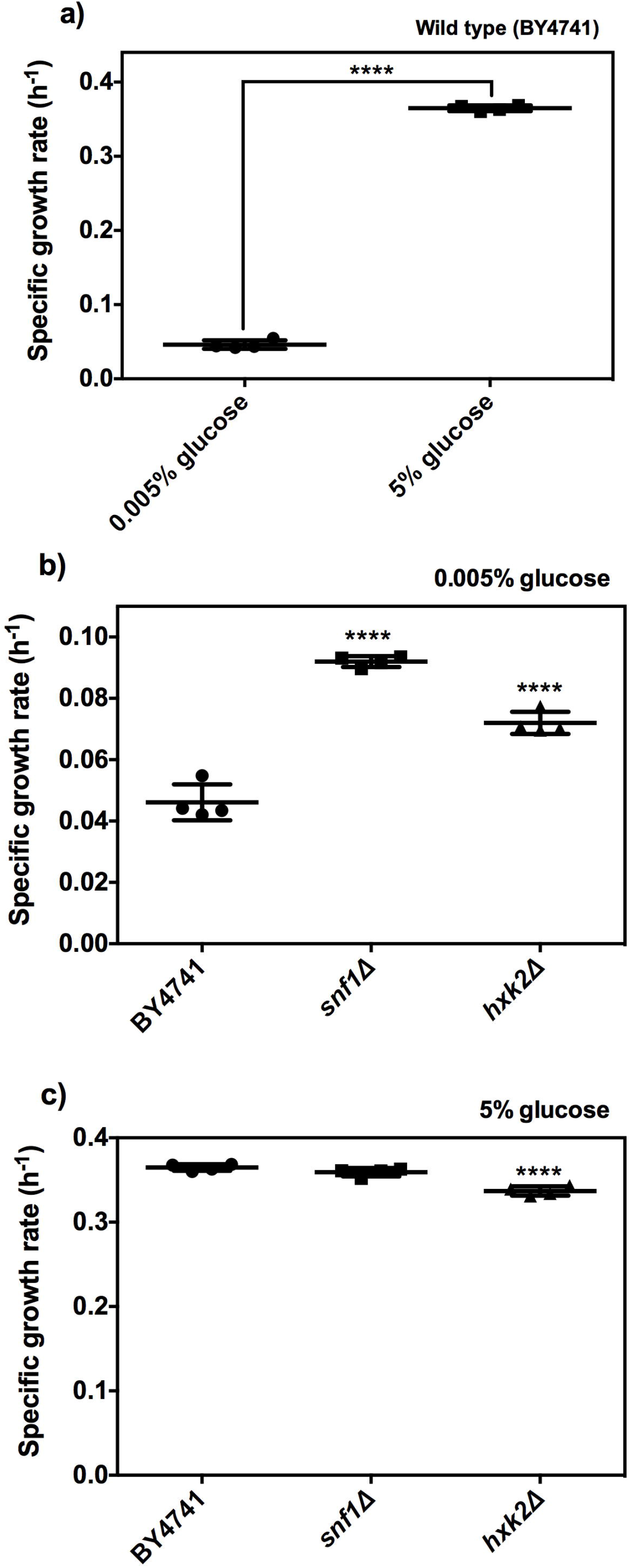
Influence of the deletion of the genes *SNF1* and *HXK2* on the exponential growth of *S. cerevisiae* at different glucose concentrations. The exponential growth was determined with the specific growth rate, which was calculated fitting data with the exponential growth rate. a) Specific growth rate of BY4741 at 0.005% and 5% glucose; b) represents growth at 0.005% glucose; c) represents growth at 5% glucose. The data represents mean ± standard deviation of four independent experiments with two technical replicates. For panels b) and c) means were compared using one-way ANOVA followed by a Dunnett multiple comparison *vs.* BY4741 (\*\*\*\**P* < 0.0001). Two-tailed unpaired *t*-test was used to compare means of the panel a) (\*\*\*\**P* < 0.0001).

### Influence of the overexpression of gene *HXK2* on the growth of *S. cerevisiae*

Hxk2p is a key modulator of the glucose repression, however, is not clear when acts as a genetic modulator and when acts as a catalytic molecule. It has been reported that in low-glucose-containing media *HXK2* expression is repressed by the transcriptional factors Rgt1p and Med8p [25, 26]. At low-glucose concentrations probably the amount of Hxk2p regulates its function. To prove this idea, we made two strains that overexpress the *HXK2* gene in the strains BY4741 and *hxk2*Δ. The gene expression was under the *GAL1* inducible promoter, which induces expression with galactose and represses expression with glucose. Besides, as a plasmid control, we used the plasmid pYES2.1 with the expression of *lacZ* gene to discard any effect produced for the extra DNA molecules within cells. Glucose supplementation of the pre-inoculum did not influence any changes in the growth of the recombinant strains BY4741//*GAL1-HXK2* and *hxk2*Δ//*GAL1-HXK2* in comparison with the parental strains BY4741 and *hxk2*Δ at 0.005% and 5% glucose (**Fig. 2a-b**). Interestingly, when galactose was used as a carbon source in the pre-inoculum to induce *HXK2* expression, the phenotype showed by the *hxk2*Δ strain at 0.005% in the recombinant strain *hxk2*Δ//*GAL1-HXK2* was reverted, decreasing its growth compared to the BY4741 strain (**Fig. 2c**). At 5% glucose, *HXK2* expression also reverts the diminution on the growth of the *hxk2*Δ strain (**Fig. 2d**). The reverting phenotype of the strain *hxk2*Δ observed by *HXK2* overexpression also corroborates that the effects on growth detected in this strain are specific of the *HXK2* deletion. Nonetheless, *HXK2* overexpression in the BY4741 strain not display any change in the growth neither at 0.005% or 5% glucose (**Fig. 2c-d**). Overall, these data indicate that overexpression of the gene *HXK2* did not have any effect on the growth of the BY4741.

**Fig 2.**
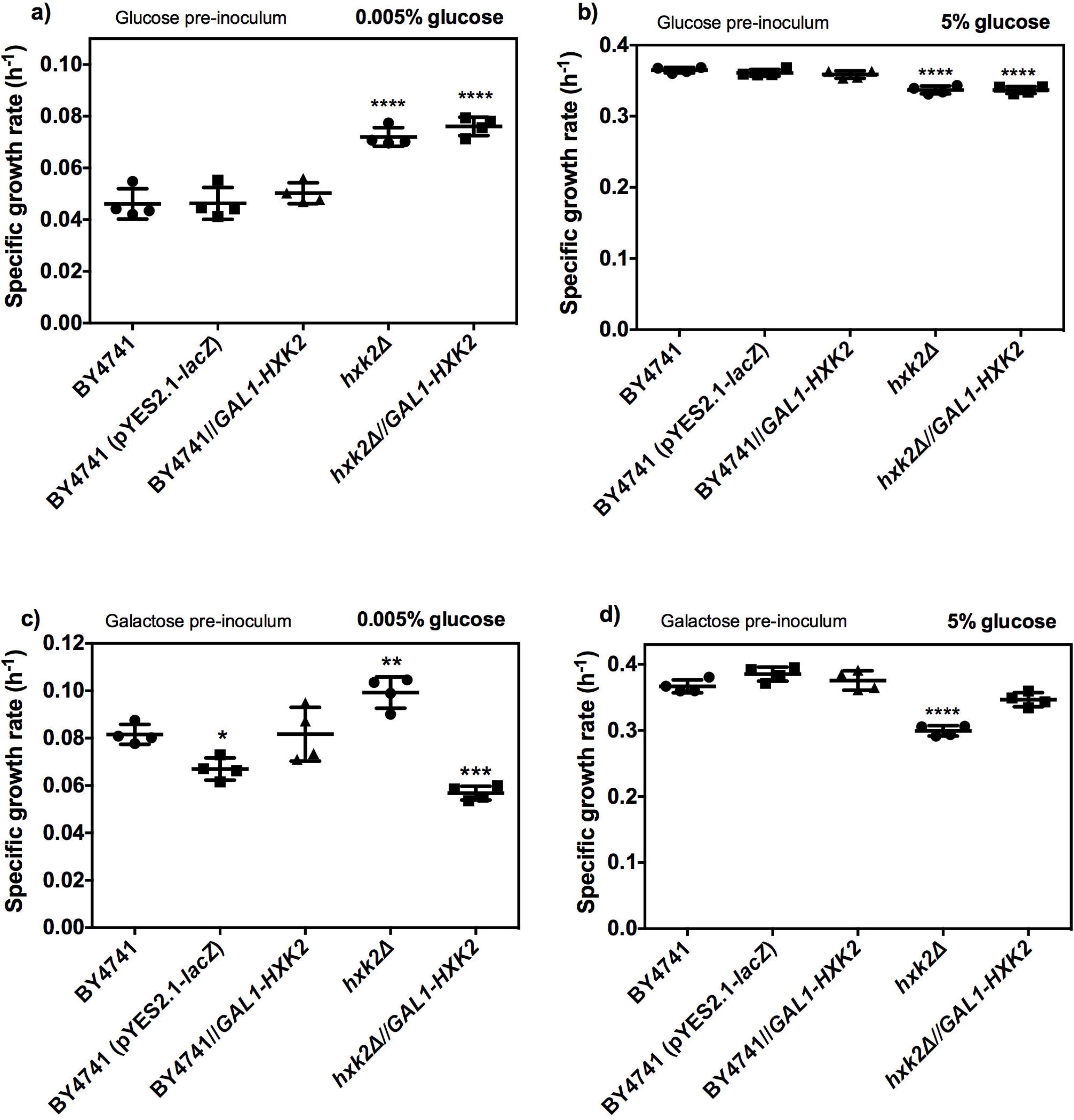
Effect of the *HXK2* gene overexpression on the exponential growth of *S. cerevisiae* strains BY4741 and *hxk2*Δ. The specific growth rate was calculated fitting data with the exponential growth rate. a) Represents growth at 0.005% glucose coming from a pre-inoculum supplemented with glucose; b) represents growth at 5% glucose coming from a pre-inoculum supplemented with glucose; c) represents growth at 0.005% glucose coming from a pre-inoculum supplemented with galactose; d) represents growth at 5% glucose coming from a pre-inoculum supplemented with galactose. The data represents mean ± standard deviation of four independent experiments with two technical replicates. Means were compared using one-way ANOVA followed by a Dunnett multiple comparison *vs.* BY4741 (\**P* < 0.05;\*\**P* < 0.01;\*\*\**P* < 0.001;\*\*\*\**P* < 0.0001).

### Impact of *HXK2* gene overexpression on *S. cerevisiae* mitochondrial respiration

Another important phenotype exerted by glucose repression is the diminution of mitochondrial respiration, which is a well-known phenotype performed by the increase of the glucose concentration in *S. cerevisiae* (**Fig. 3a**). Therefore, the measuring of the mitochondrial respiration is a valuable indicator of glucose repression that could help to obtain further comprehension into the role of the *SNF1* and *HXK2* genes in the glucose repression. The yeast cultures utilized to mitochondrial respiration assay came from a pre-inoculum supplemented with galactose to ensure the *HXK2* expression.

**Fig 3.**
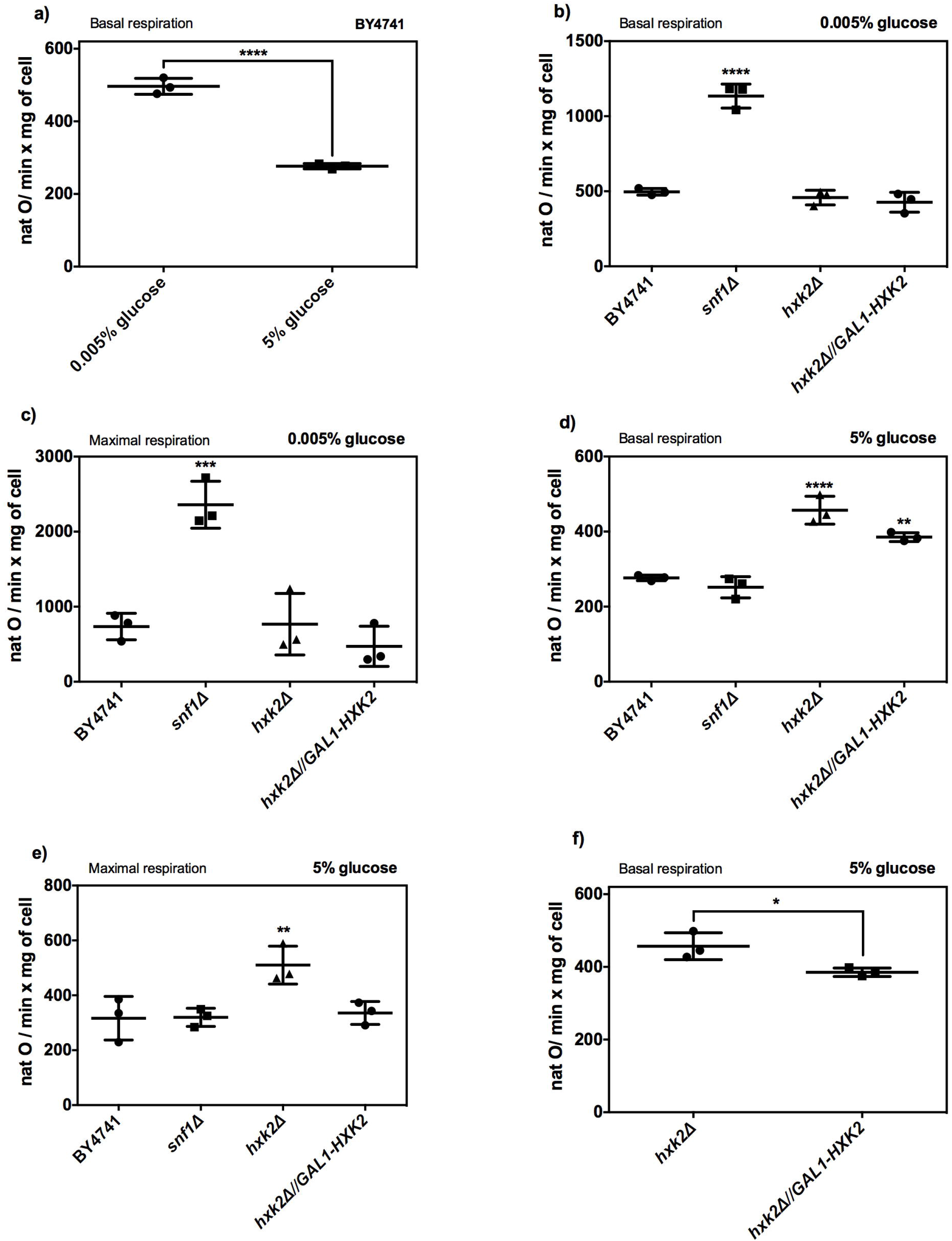
Mitochondrial respiration of the strains BY4741, *hxk2*Δ, *snf1*Δ, and *hxk2*Δ//*GAL1-HXK2* at 0.005% and 5% glucose. Basal and maximal respiration were obtained from cultures in which pre-inoculum was grown with galactose in an SC medium. a) Basal respiration of BY4741 strain at 0.005% and 5% glucose; b) basal respiration of the strains BY4741, *hxk2*Δ, *snf1*Δ, and *hxk2*Δ//*GAL1-HXK2* at 0.005% glucose; c) maximal respiration of the strains BY4741, *hxk2*Δ, *snf1*Δ, and *hxk2*Δ//*GAL1-HXK2* at 0.005% glucose; d) basal respiration of the strains BY4741, *hxk2*Δ, *snf1*Δ, and *hxk2*Δ//*GAL1-HXK2* at 5% glucose; e) maximal respiration of the strains BY4741, *hxk2*Δ, *snf1*Δ, and *hxk2*Δ//*GAL1-HXK2* at 5% glucose; f) comparison of the basal respiration between *hxk2*Δ and *hxk2*Δ//*GAL1-HXK2* strains at 5% glucose. The data represents mean ± standard deviation of three independent experiments. For panels b), c), d), and e) means were compared using one-way ANOVA followed by a Dunnett multiple comparison *vs.* BY4741 (\*\**P* < 0.01;\*\*\**P* < 0.001;\*\*\*\**P* < 0.0001). For panels a) and f) means were compared with a two-tailed unpaired *t*-test (\**P* < 0.05; \*\*\**P* < 0.001).

Interestingly, under the low concentration glucose (0.005%) the *snf1*Δ strain increased the basal and maximal mitochondrial respiration in contrast with the BY4741 strain (**Fig. 3b-c**). On the contrary, the strains *hxk2*Δ and *hxk2*Δ//*GAL1-HXK2* did not show differences in basal and maximal mitochondrial respiration (**Fig. 3b-c**). At 5% glucose the strains *hxk2*Δ and *hxk2*Δ//*GAL1-HXK2* increased the basal respiration compared to the BY4741 strain (**Fig. 3d**); while only the strain *hxk2*Δ increased the maximal respiration (**Fig. 3e**). Importantly, between strains *hxk2*Δ and *hxk2*Δ//*GAL1-HXK2* exists a difference in basal respiration with a diminution of oxygen consumption by *hxk2*Δ//*GAL1-HXK2* strain (**Fig. 3f**). These data suggest that an increase of mitochondrial respiration may have a relation with the heightened in the growth in *snf1*Δ strain at 0.005% glucose. Besides, the *hxk2*Δ complementation with *HXK2* gene does not have any effect in mitochondrial respiration at 0.005% glucose, whereas at 5% glucose a decrease on mitochondrial respiration is appreciated due to the *HXK2* expression. Probably the quantity of *HXK2* expression is necessary to repress mitochondrial respiration in high-glucose concentrations.

### Influence of the deletion of the genes *HXK2,* and *SNF1* on the transcription of *S. cerevisiae* BY4741

To further understand whether the *SNF1/HXK2* pathway modulates the growth and mitochondrial respiration through hexose transporters genetic regulation, we decide to measure the transcription of the hexoses transporters that allow us to recognize the interaction among these phenotypes and the pathway *SNF1/HXK2*. Yeast cultures of these experiments came from a pre-inoculum grown with galactose to induce the expression in the recombinant strain *hxk2*Δ//*GAL1-HXK2* and were grown in SC minimal media.

According to its affinity, we choose to quantify the transcription of the low-affinity hexose transporters genes (*HXT1* and *HXT3*) and the high-affinity hexose transporters genes (*HXT2* and *HXT4*). In the first place, the transcription in the strain BY4741 was used to corroborate the change in the transcription of *HXT1-4* genes in relation to the glucose concentration. *HXT1* and *HXT3* transcription augment at 5% of glucose in comparison with the yeast grown with 0.005% glucose (**Fig 4a-b**). On the contrary, transcription of the gene *HXT2* decreases at 5% glucose with respect to the cultures grown at 0.005% glucose (**Fig 4c**). Although, *HXT4* has been classified as a transporter of moderate or high affinity did not show any change in its transcription levels (**Fig 4d**). These data corroborate by an absolute quantification that *HXT1* and *HXT3* transcription is accord to its classification as low-affinity hexose transporters, likewise *HXT2* gene transcript induction, that occurs under low glucose concentration, corresponding to a high-affinity hexose transporter. However, *HXT4* transcription did not change between the glucose concentrations evaluated.

**Fig 4.**
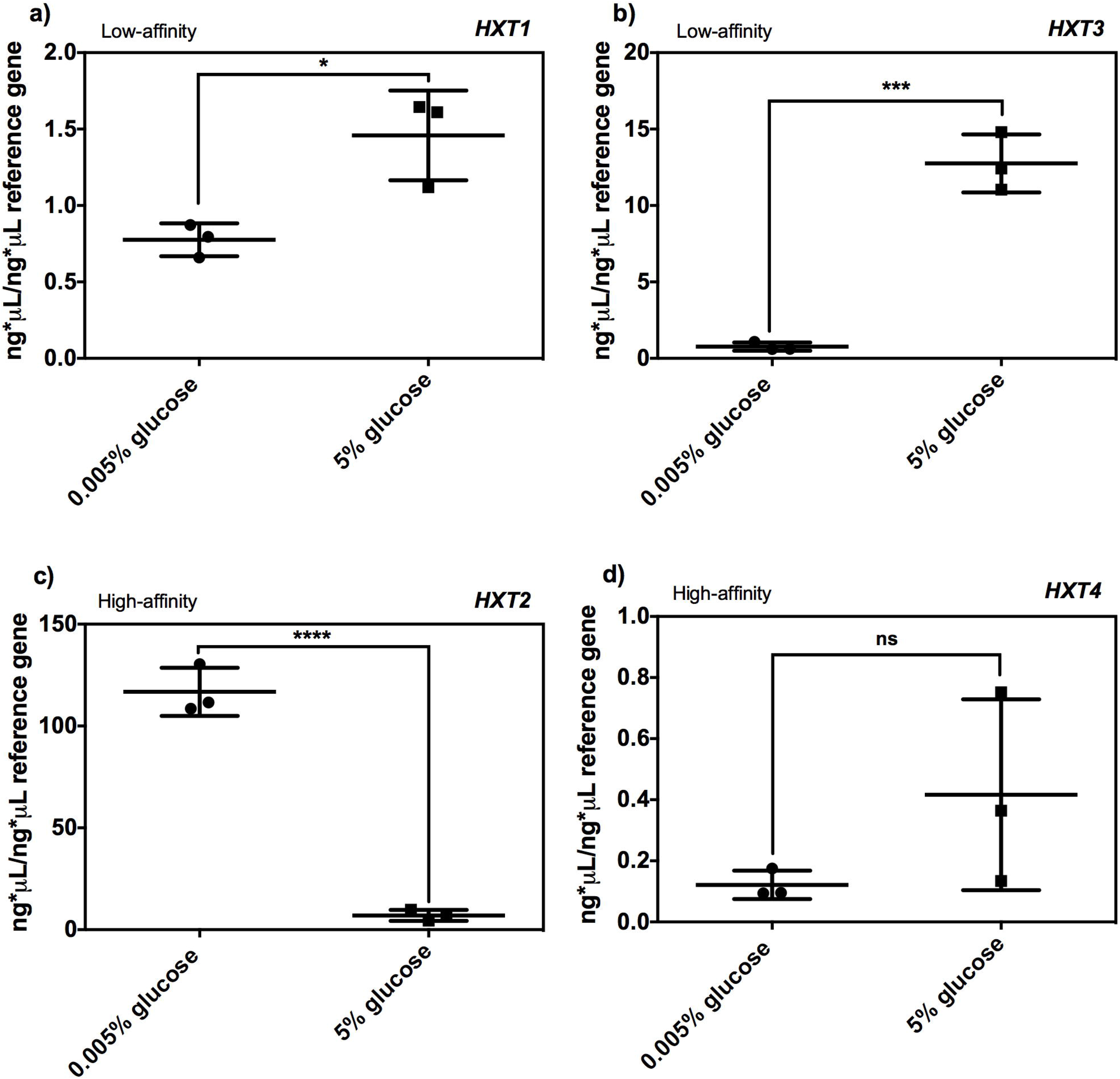
Transcription of the genes *HXT1*, *HXT2*, *HXT3*, and *HXT4* in the BY4741 strain at 0.005% and 5% glucose. Absolute quantification was made using a *UBC6* as a reference gene, from *S. cerevisiae* cultures in which pre-inoculum was grown with galactose in an SC medium. a) Transcription of *HXT1* gene; b) transcription of *HXT3* gene; c) Transcription of *HXT2* gene; d) transcription of *HXT4* gene. The data represents mean ± standard deviation of three independent experiments with two technical replicates. Means were compared with a two-tailed unpaired *t*-test (\**P* < 0.05; \*\*\**P* < 0.001; \*\*\*\**P* < 0.0001; ns, non-significant).

Low-affinity hexose transporters like *HXT1* and *HXT3* are expressed at high-glucose concentrations mainly. At high glucose concentrations, Snf1p is mainly inactive; *snf1*Δ strain could resemble an inactive phenotype of this protein. Deletion of the gene *SNF1* increases *HXT1* transcription at 5% glucose and does not have any effect in its transcription at 0.005% glucose (**Fig. 5a-b**); meanwhile, *HXT3* transcription increased in the *snf1*Δ strain at 0.005% and did not display differences in transcription at 5% glucose (**Fig. 5c-d**). In the case of the *HXK2* gene deletion and its complementing strain *hxk2*Δ//*GAL1-HXK2* both showed a decrease in the transcription levels of the genes *HXT1* and *HXT3* at 5% glucose and at 0.005% glucose only *HXT1* transcription showed a decrease (**Fig. 5**). Under low concentrations of glucose (>0.1%) high-affinity transporters *HXT2* and *HXT4* are maximally expressed. Transcription of the *HXT4* gene increased in the *snf1*Δ strain at 0.005% and 5% glucose, whereas *HXT2* transcription was unaffected by *snf1*Δ strain (**Fig. 6**). *HXK2* deletion enhanced transcription of the genes *HXT2* and *HXT4* only at 5% glucose (**Fig. 6**). Importantly, overexpression of the gene *HXK2* reverted the increase in transcriptional levels of the genes *HXT2* and *HXT4* showed by the *hxk2*Δ strain at 5% glucose (**Fig. 6**). In sum, these results suggest a negatively transcriptional regulation of *HXK2* upon high-affinity transporters *HXT2* and *HXT4*, since overexpression of *HXK2* maintains the transcription at wild-type levels. However, transcription of low-affinity transporters has repressed, even in the *hxk2*Δ//*GAL1-HXK2* strain. Nonetheless, *SNF1* deletion did not showed a pattern in transcription regulation, increasing *HXT1* and *HXT4* transcription at 5% glucose and *HXT2* and *HXT4* at 0.005% glucose.

**Fig 5.**
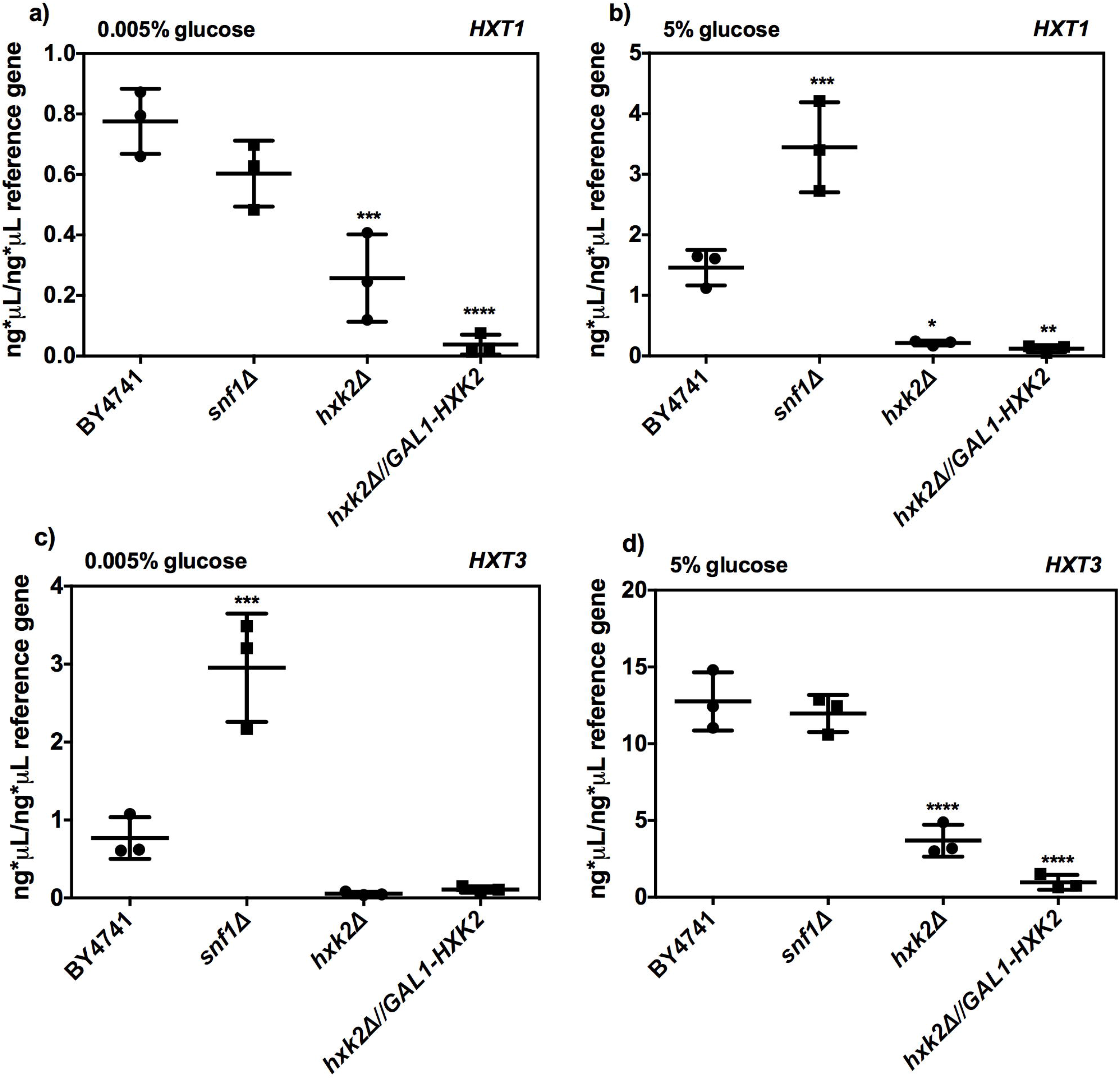
Transcription of the low-affinity hexose transporters genes *HXT1* and *HXT3* in the BY4741, *hxk2*Δ, *snf1*Δ, and *hxk2*Δ//*GAL1-HXK2* strains at 0.005% and 5% glucose. Absolute quantification was made using a *UBC6* as a reference gene, from *S. cerevisiae* cultures in which pre-inoculum was grown with galactose in an SC medium. a) Represents *HXT1* gene transcription at 0.005%; b) represents *HXT1* gene transcription at 5%; c) represents *HXT3* gene transcription at 0.005%; b) represents *HXT3* gene transcription at 5%. The data represents mean ± standard deviation of three independent experiments. Means were compared using one-way ANOVA followed by a Dunnett multiple comparison *vs.* BY4741 (\**P* < 0.05; \*\**P* < 0.01; \*\*\**P* < 0.001; \*\*\*\**P* < 0.0001).

**Fig 6.**
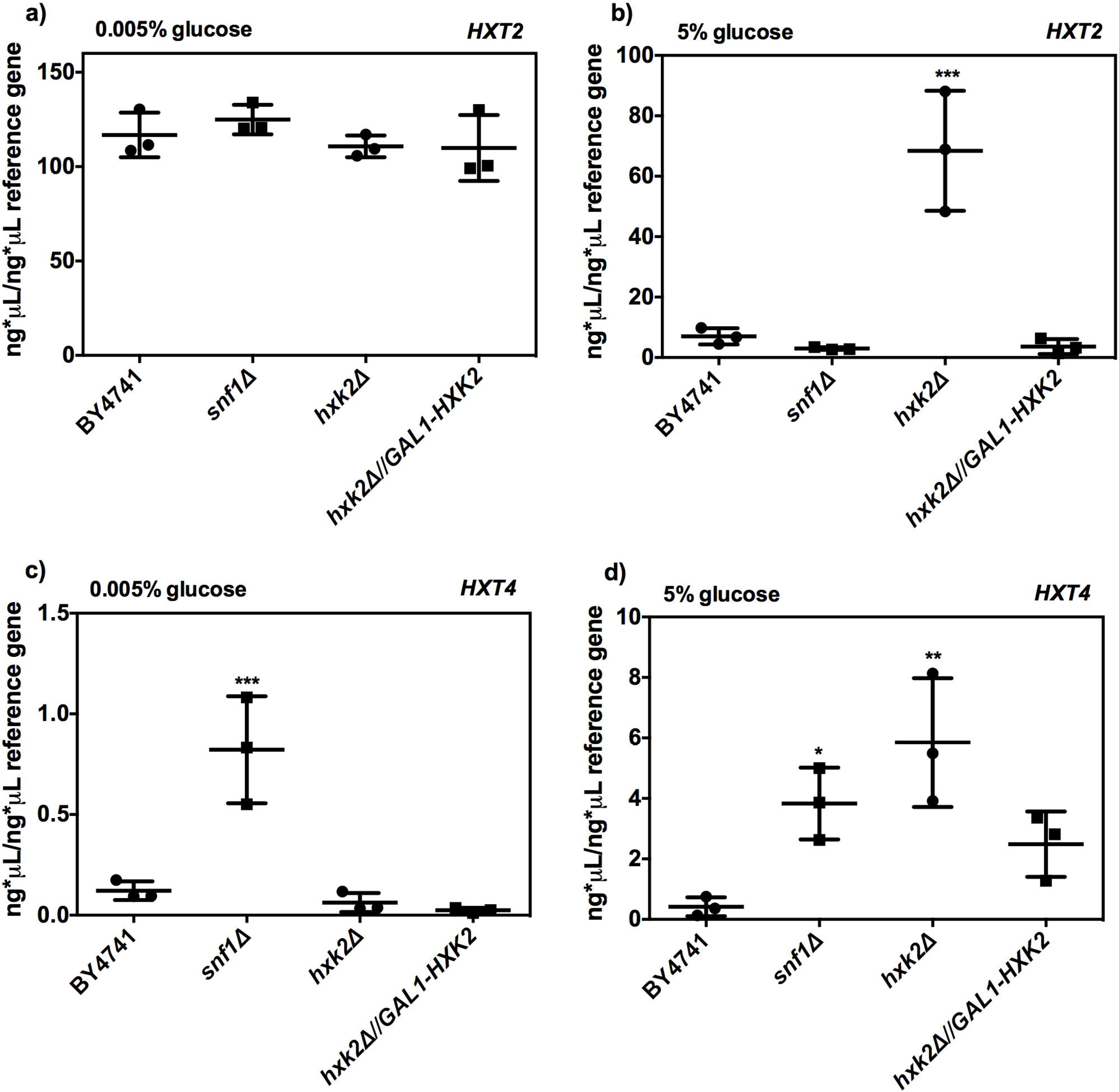
Transcription of the high-affinity hexose transporters genes *HXT2* and *HXT4* in the BY4741, *hxk2*Δ, *snf1*Δ, and *hxk2*Δ//*GAL1-HXK2* strains at 0.005% and 5% glucose. Absolute quantification was made using a *UBC6* as a reference gene, from *S. cerevisiae* cultures in which pre-inoculum was grown with galactose in an SC medium. a) Represents *HXT2* gene transcription at 0.005%; b) represents *HXT2* gene transcription at 5%; c) represents *HXT4* gene transcription at 0.005%; b) represents *HXT4* gene transcription at 5%. The data represents mean ± standard deviation of three independent experiments. Means were compared using one-way ANOVA followed by a Dunnett multiple comparison *vs.* BY4741 (\**P* < 0.05; \*\**P* < 0.01; \*\*\**P* < 0.001).

### The relationship among exponential growth, mitochondrial respiration, and hexose transporters transcription

A correlational analysis was carried out to discriminate whether exists an effect of hexose transporters upon exponential growth and mitochondrial respiration in *S. cerevisiae*. The biochemical parameter *Vmax* was chosen to compare the relation of the hexose transporters with exponential growth and mitochondrial respiration, due to represents the velocity of glucose transport, which may impact the glycolytic flux. *Vmax* values were obtained from Reifenberger, Boles and Ciriacy [6]. Hexose transporters mainly transcribed, exponential growth values, and mitochondrial respiration were summarized in **table 2** for 0.005% glucose, and **table 3** for 5% glucose. For the correlational analysis, we only plotted the *Vmax* values of the hexose transporter genes with high transcriptional levels, in the case that were two genes with an increase of the transcriptional levels the average of them was plotted. We found a positive correlation between exponential growth and *Vmax* (r = 0.8452; *P* = 0.0082; **Fig. 7a**); indicating those hexose transporters with the major *Vmax* have the greater exponential growth, corresponding to the yeast cultures grown at 5% glucose, except for *hxk2*Δ. The mitochondrial respiration showed a negative correlation with the *Vmax* values (r = −0.8201; *P* = 0.0239; **Fig. 7b**); this data suggest that hexose transporters with low-values of *Vmax* promote mitochondrial respiration and hexose transporters with high-values of *Vmax* decrease mitochondrial respiration. Entirely, these data suggest a strong relation between glucose transporters with the exponential growth and mitochondrial respiration.

**Table 2.**
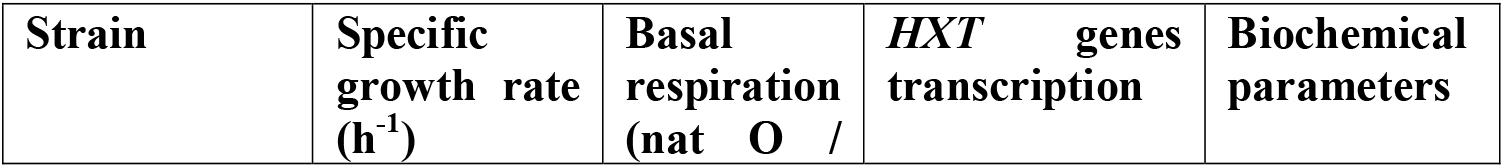
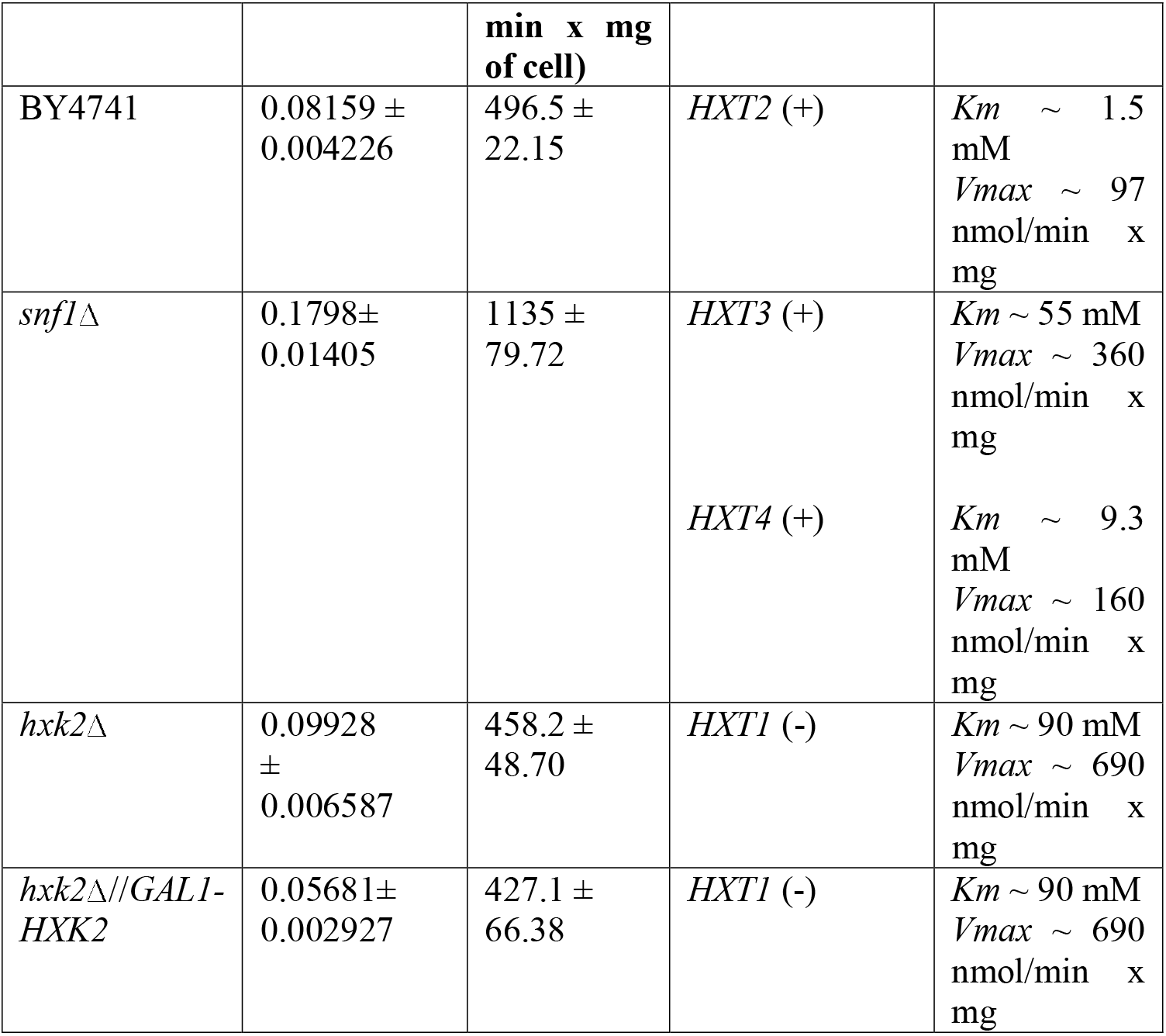
Summary of phenotypes showed by BY4741, *snf1*Δ, *hxk2*Δ, and *hxk2*Δ//*GAL1-HXK2* at 0.005% glucose.

**Table 3.**
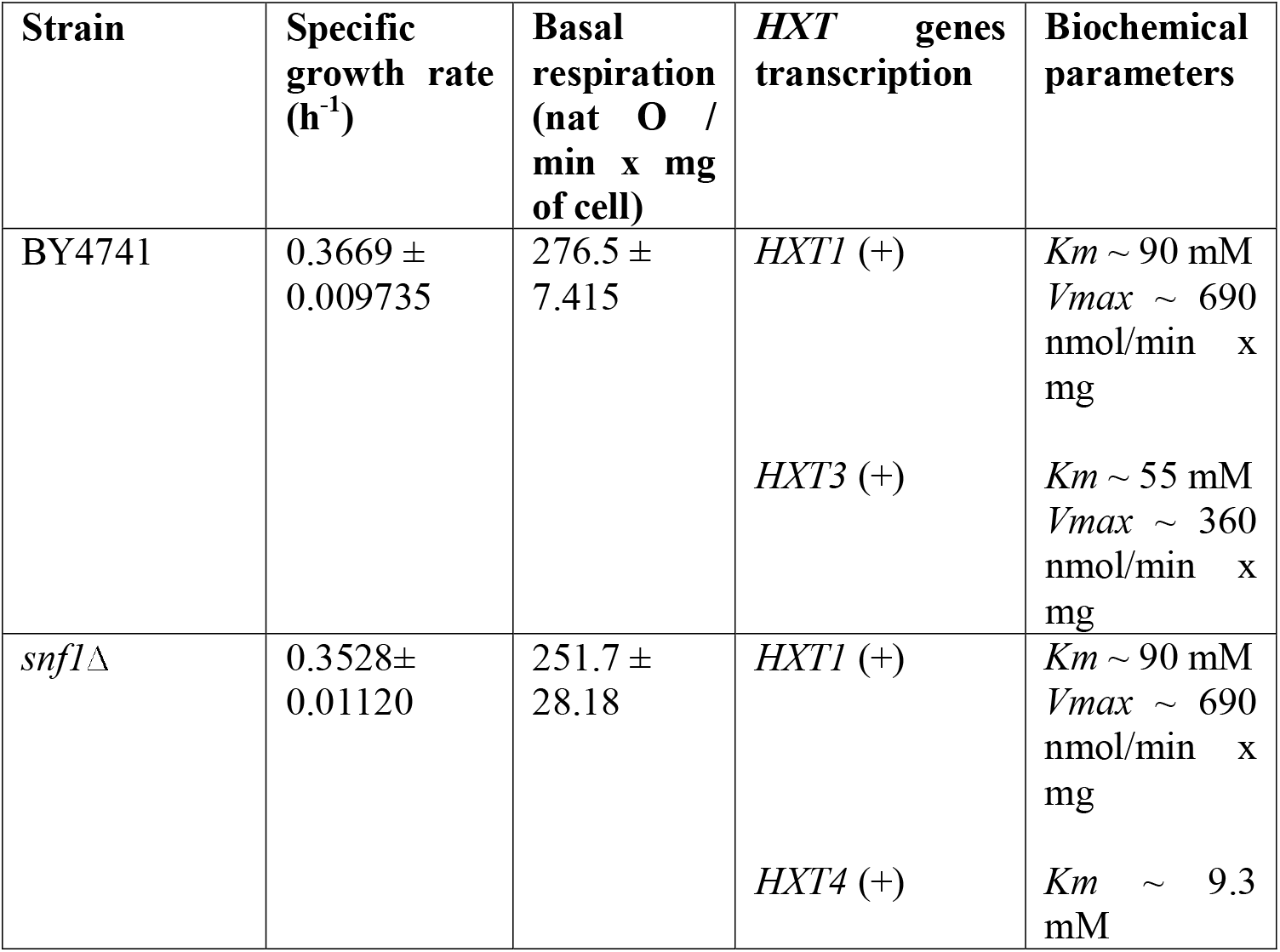
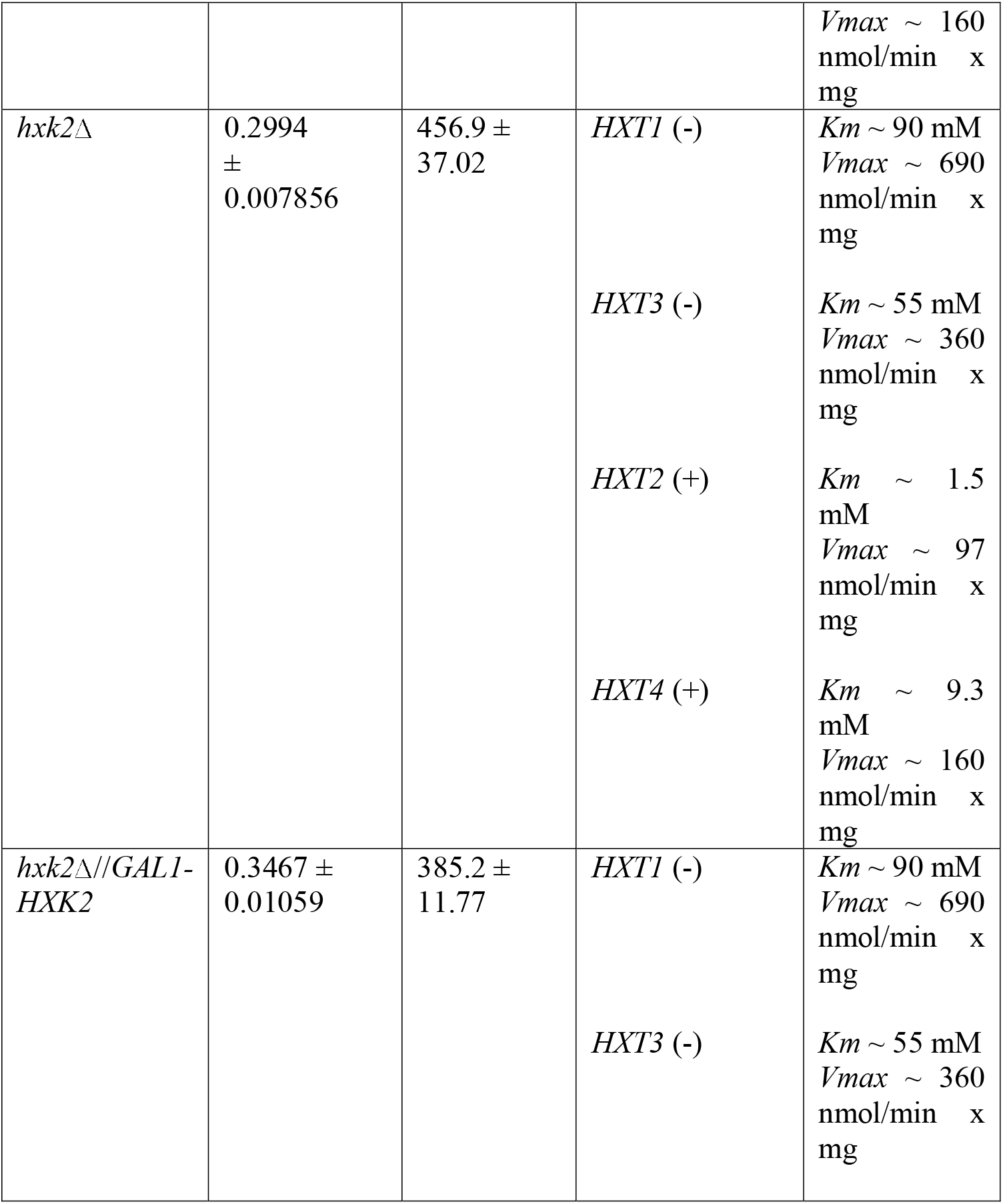
Summary of phenotypes showed by BY4741, *snf1*Δ, *hxk2*Δ, and *hxk2*Δ//*GAL1-HXK2* at 5% glucose.

**Fig 7.**
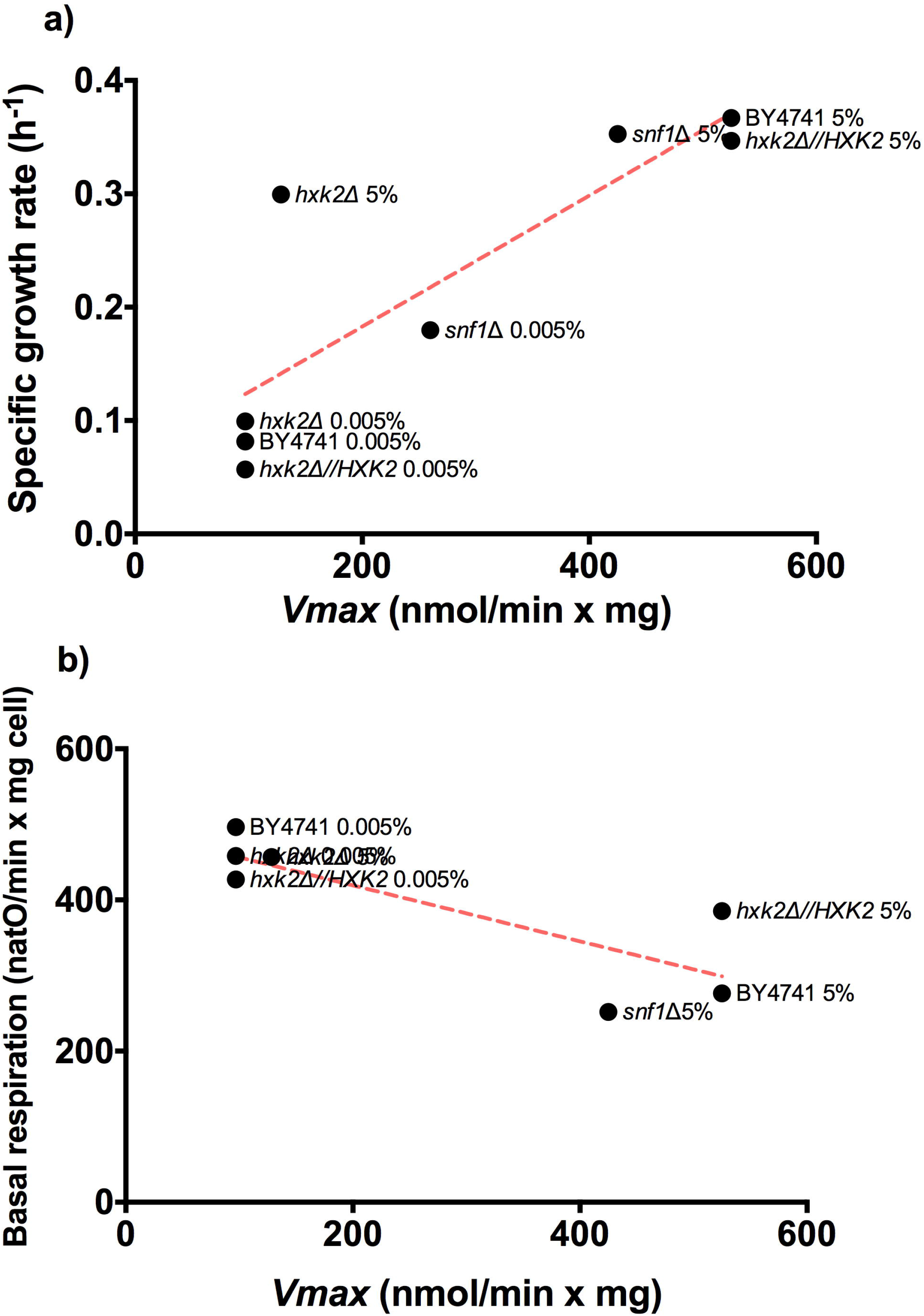
Analysis of correlation between exponential growth/mitochondrial respiration with *Vmax* of hexose transporters. Pearson’s correlational analysis was carried out to evaluated the correlation between exponential growth/mitochondrial respiration with *Vmax*. a) Exponential growth *vs. Vmax*; b) mitochondrial respiration *vs. Vmax*. Pearson’s correlation was calculated with the statistical package GraphPad Prism 6.00 for Macintosh (GraphPad Software).

### Effect of the deletion of the genes *MIG1* and *SNF1* on the transcription of *S. cerevisiae* BY4741

To a better understanding of the relation between the *SNF1* pathway and transcriptional levels of hexose transporters, we decided to make an additional experiment using a different *S. cerevisiae* strain BY4742 and its mutant in the *SNF1* gene, to discard strain-specific effects. Additionally, we monitored the transcription of two additional hexose transporters *HXT6* and *HXT7*, which are classified as high-affinity transporters. Besides, we use 0.01% (0.55 mM) as a low glucose concentration, to take a wider range of glucose concentrations. Finally, these experiments were carried out in YPD medium and came from a pre-inoculum grown with glucose.

To corroborate the expression of the hexose transporters according to its classification, we decided to measure its transcription under low glucose concentration (0.01%) and high glucose concentration (5%). Low-affinity transporters (*HXT1* and *HXT3*) increased its transcription at 5% glucose in comparison with transcription at 0.01% glucose (**Fig. 8a-b**). In the case of *HXT2* and *HXT4* transporters are considered as moderate-affinity or high-affinity transporters, we found that *HXT2* augmented its transcription under low glucose concentration (0.01%) as high-affinity transporters do, while *HXT4* enhanced its transcription at 5% glucose like a low-affinity transporter (**Fig. 8c-d**). Finally, the high-affinity transporters *HXT6* and *HXT7* did not display any change in its transcription with the two concentrations of glucose assayed (**Fig. 8e-f**). The profile of hexose transporter transcription is the same of the presented with BY4741 strain. However, the *HXT4* presents a low-affinity transcription instead of a high-affinity pattern of transcription.

**Fig 8.**
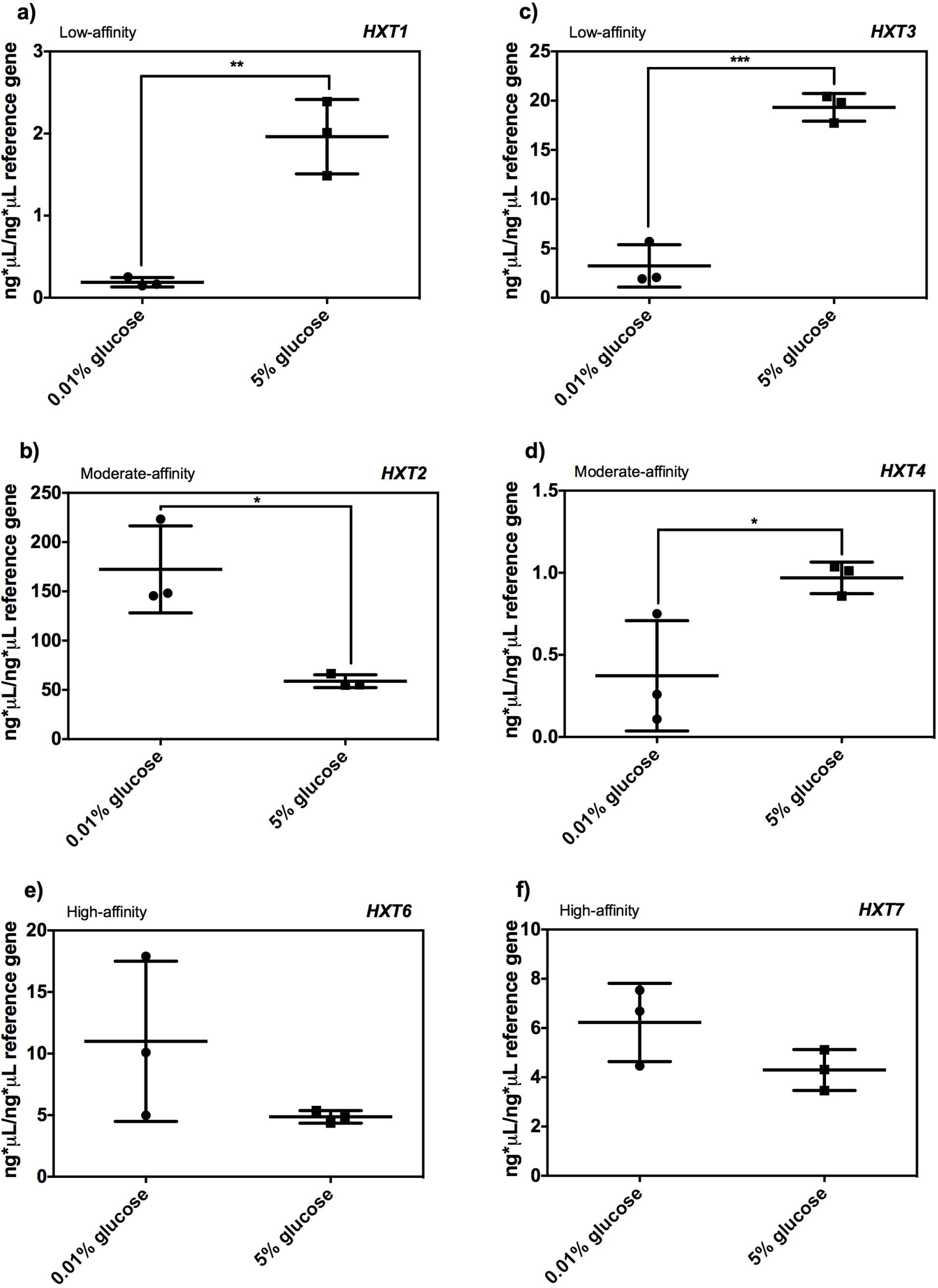
Transcription of the genes *HXT1*, *HXT2*, *HXT3*, *HXT4, HXT6,* and *HXT7* in the BY4742 strain at 0.01% and 5% glucose. Absolute quantification was made using a *UBC6* as a reference gene, from *S. cerevisiae* cultures in which pre-inoculum was grown with glucose in a YPD medium. a) Transcription of *HXT1* gene; b) transcription of *HXT3* gene; c) Transcription of *HXT2* gene; d) transcription of *HXT4* gene; e) transcription of *HXT6* gene; f) transcription of *HXT7* gene. The data represents mean ± standard deviation of three independent experiments with two technical replicates. Means were compared with a two-tailed unpaired *t*-test (\**P* < 0.05; \*\**P* < 0.01; \*\*\**P* < 0.001).

*SNF1* deletion diminished the transcription of the *HXT1* gene at 0.01% and did not change its transcription at 5% glucose (**Fig. 9a**). *HXT2* transcription was down at 0.01% and 5% glucose in the *snf1*Δ strain (**Fig. 9c**). Transcription of the *HXT6* gene was decreased in *snf1*Δ strain grown at 5% glucose with respect to BY4742 strain (**Fig. 9e**). A dual effect was observed in the transcription of the gene *HXT7*, at 0.01% *SNF1* deletion increased 10 times its transcription, whereas, at 5% *SNF1* deletion lessened *HXT7* transcription (**Fig. 9f**). Transcription of the genes *HXT3* and *HXT4* passed unaltered by the *SNF1* gene deletion (**Fig. 9b-d**).

**Fig 9.**
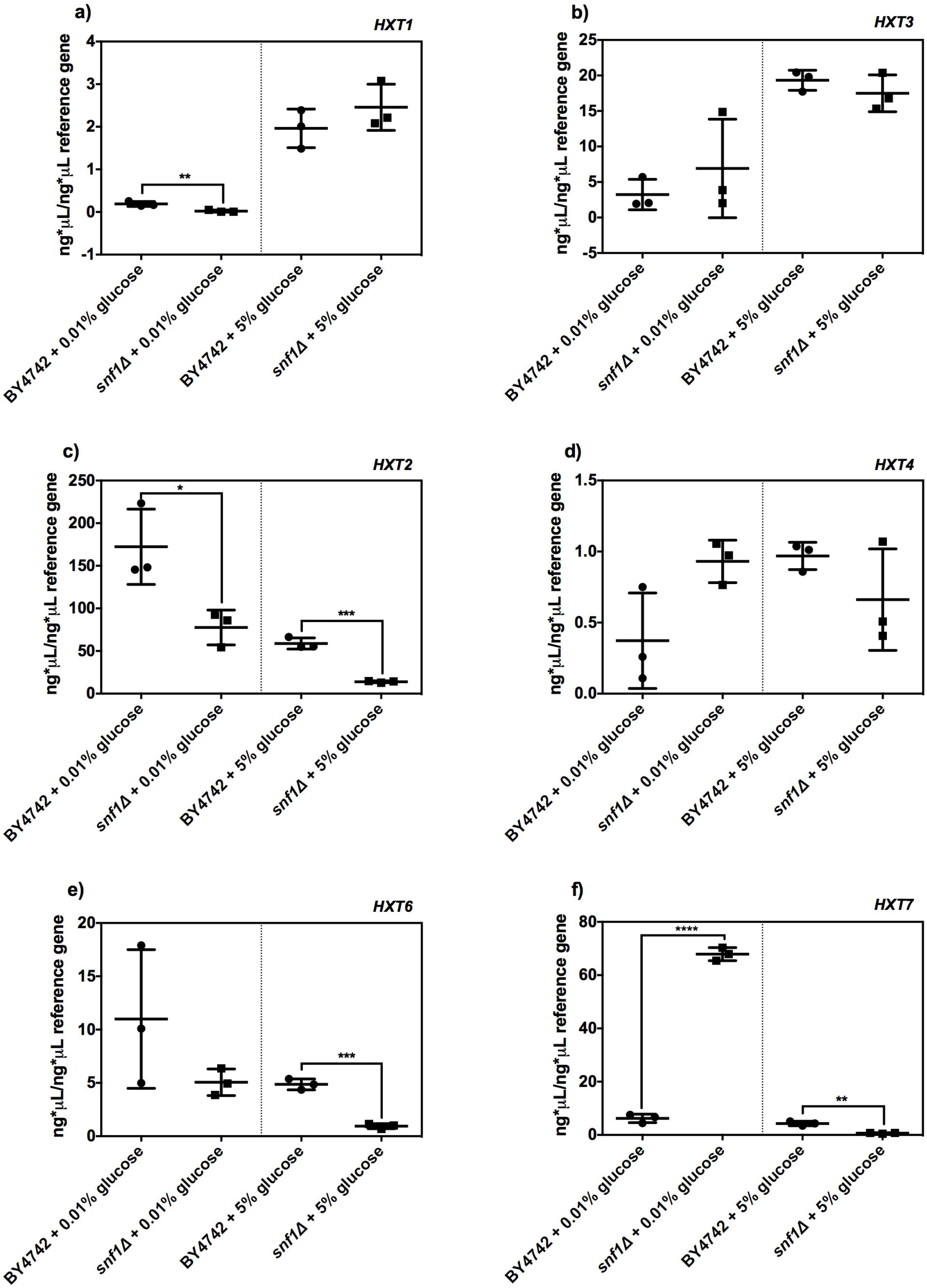
Transcription of the genes *HXT1*, *HXT2*, *HXT3*, *HXT4, HXT6,* and *HXT7* in the BY4742 and *snf1*Δ strains at 0.01% and 5% glucose. Absolute quantification was made using a *UBC6* as a reference gene, from *S. cerevisiae* cultures in which pre-inoculum was grown with glucose in a YPD medium. a) Transcription of *HXT1* gene; b) transcription of *HXT3* gene; c) Transcription of *HXT2* gene; d) transcription of *HXT4* gene; e) transcription of *HXT6* gene; f) transcription of *HXT7* gene. The data represents mean ± standard deviation of three independent experiments with two technical replicates. Means were compared with a two-tailed unpaired *t*-test (\**P* < 0.05; \*\**P* < 0.01; \*\*\**P* < 0.001).

To strength the analysis of correlation between exponential growth and *Vmax*, we included the data obtained with the BY4742 strain and its deleting strain in *SNF1*. Exponential growth data and hexose transcription were summarized in **table 4** for 0.01% glucose and **table 5** for 5% glucose. The positive correlation between exponential growth and *Vmax* was maintained (r = 0.8495; *P* = 0.0005; **Fig. 10**); showing a relation between the exponential growth at 5% glucose concentration with the higher values of *Vmax*, and exponential growth at 0.005% and 0.01% glucose with the lower values of *Vmax.* This result suggests that *Vmax* of the hexose transporters could regulate the exponential growth of *S. cerevisiae*.

**Table 4.** Summary of phenotypes showed by BY4742 and *snf1* at 0.01% glucose.

| Strain | Specific growth rate (h<sup>-1</sup>) | *HXT* gene transcription augmented | Biochemical parameter of *HXT* most transcribed |  |
| --- | --- | --- | --- | --- |
| BY4742 | 0.1118 ±<br>0.008450 | *HXT2* (+) | $K_m \sim 1.5$ mM<br>$V_{max} \sim 97$ nmol/min x mg | |
| *snf1Δ* | 0.02867 ±<br>0.005741 | *HXT7* (+) | $K_m \sim 1.1$ mM<br>$V_{max} \sim 186$ nmol/min x | |
|  |  | *HXT1* (-) | mg<br>*Km* ~ 90 mM<br>*Vmax* ~ 690<br>nmol/min x<br>mg |  |
|  |  | *HXT2* (-) | *Km* ~ 1.5<br>mM<br>*Vmax* ~ 97<br>nmol/min x<br>mg |  |

**Table 5.** Summary of phenotypes showed by BY4742 and *snf1* at 5% glucose.

| Strain | Specific growth rate (h<sup>-1</sup>) | *HXT* gene transcription augmented | Biochemical parameter of *HXT* most transcribed | 900 |
| --- | --- | --- | --- | --- |
| BY4742 | 0.3255 ± 0.008632 | *HXT1* (+)<br>*HXT3* (+) | *Km* ~ 90 mM<br>*Vmax* ~ 690<br>nmol/min x<br>mg<br>*Km* ~ 55 mM<br>*Vmax* ~ 360<br>nmol/min x<br>mg |  |
| *snf1*Δ | 0.3433 ± 0.01653 | *HXT2* (-)<br>*HXT6* (-)<br>*HXT7* (-) | *Km* ~ 1.5<br>mM<br>*Vmax* ~ 97<br>nmol/min x<br>mg<br>ND<br>*Km* ~ 1.1<br>mM<br>*Vmax* ~ 186<br>nmol/min x<br>mg |  |

**Fig 10.**
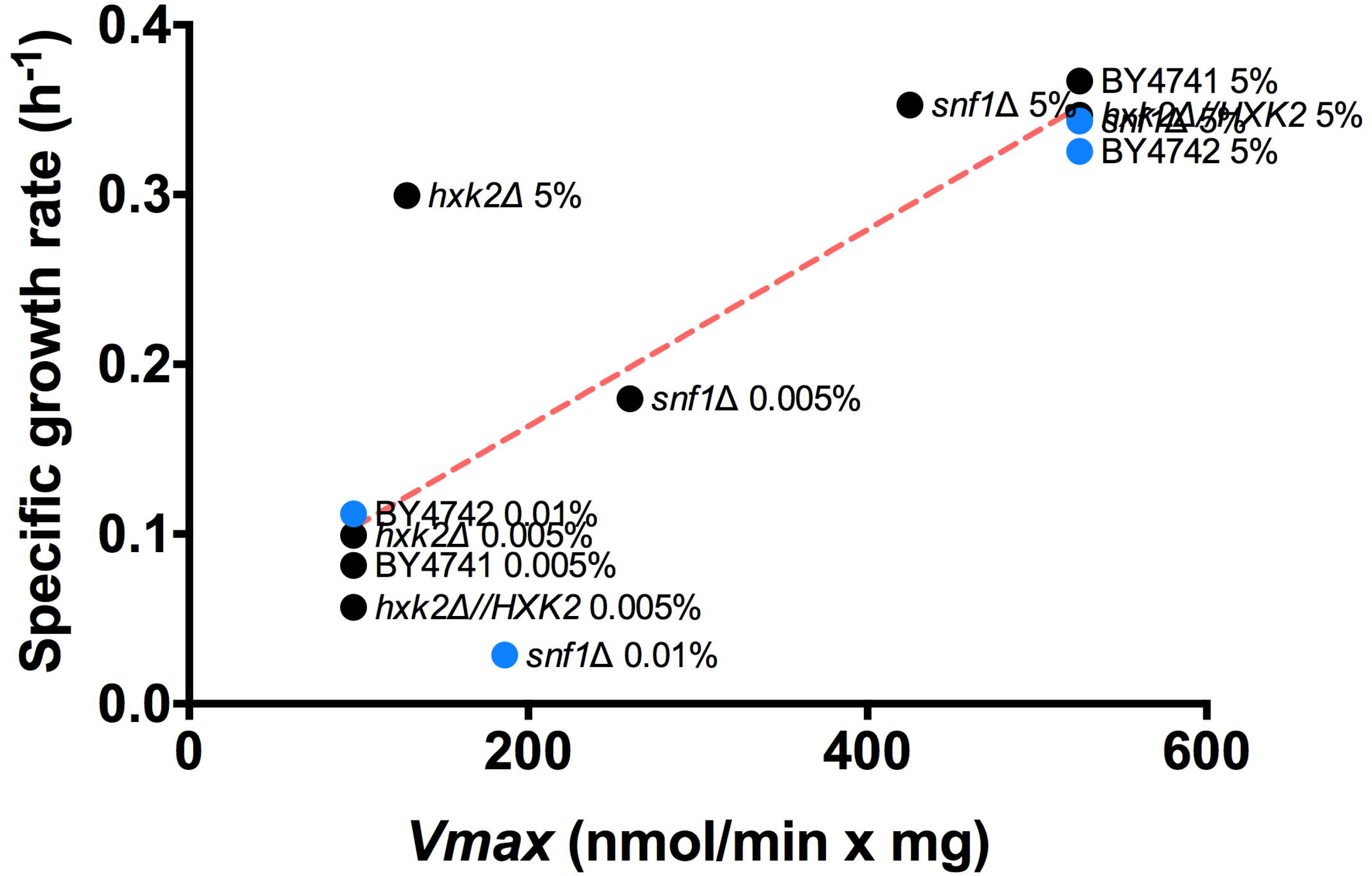
Analysis of correlation between exponential growth with *Vmax* of hexose transporters. Pearson’s correlational analysis was carried out to evaluated the correlation between exponential growth with *Vmax*. Black dots represent *S. cerevisiae* cultures in which pre-inoculum was grown with galactose in an SC medium. Blue dots represent *S. cerevisiae* cultures in which pre-inoculum was grown with glucose in a YPD medium. Pearson’s correlation was calculated with the statistical package GraphPad Prism 6.00 for Macintosh (GraphPad Software).

To gain knowledge about the hexose transporters and its regulation by the Snf1p pathway, we decided to make a multivariate data analysis that included the transcriptional level data form hexose transporters obtained from strains BY4741 and BY4742 and its deleting strains in *SNF1* gene. Importantly, PCA analysis showed that more dispersed clustering of hexose transcription corresponds to the moderate-affinity transporters especially the *HXT2* transcription (**Fig. 11**). *HXT2* transcription was enhanced by low-glucose concentrations in both strains measured. At 5% glucose *HXK2* deletion increased *HXT2* transcription, while *SNF1* deletion decreased it.

**Fig 11.**
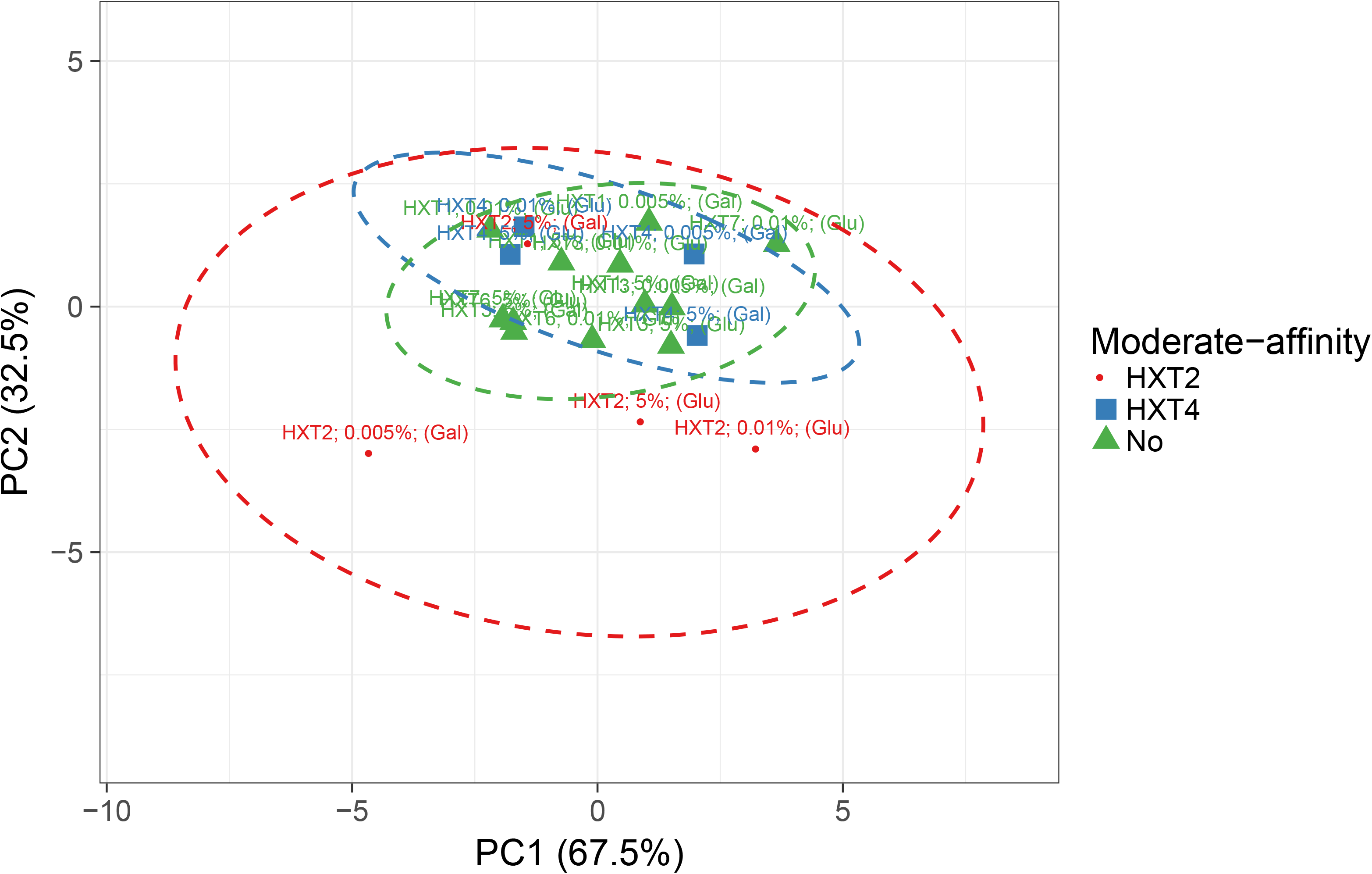
PCA analysis of *HXT* transcription in the strain BY4741 and BY4742, and its strains deleted in the *SNF1* gene. Original values are ln (x + 1)-transformed. No scaling is applied to rows; probabilistic PCA is used to calculate principal components. *X* and *Y* axis show principal component 1 and principal component 2 that explains 67.5% and 32.5% of the total variance, respectively. Prediction ellipses are such that with probability 0.95, a new observation from the same group will fall inside the ellipse. N = 20 data points.

To corroborate the effect of the Snf1p/Hxk2p pathway upon *HXT2* transcription was used *MIG1* as an additional gene involved in the pathway. As expected, the deletion of the *MIG1* gene had a dual effect upon *HXT2* transcription, increasing its transcription at 5% glucose and decreasing it at 0.01% glucose (**Fig. 12**). These data indicate that the *SNF1/HXK2/MIG1* pathway might have a specific role in the regulation of the *HXT2* gene transcription.

**Fig 12.**
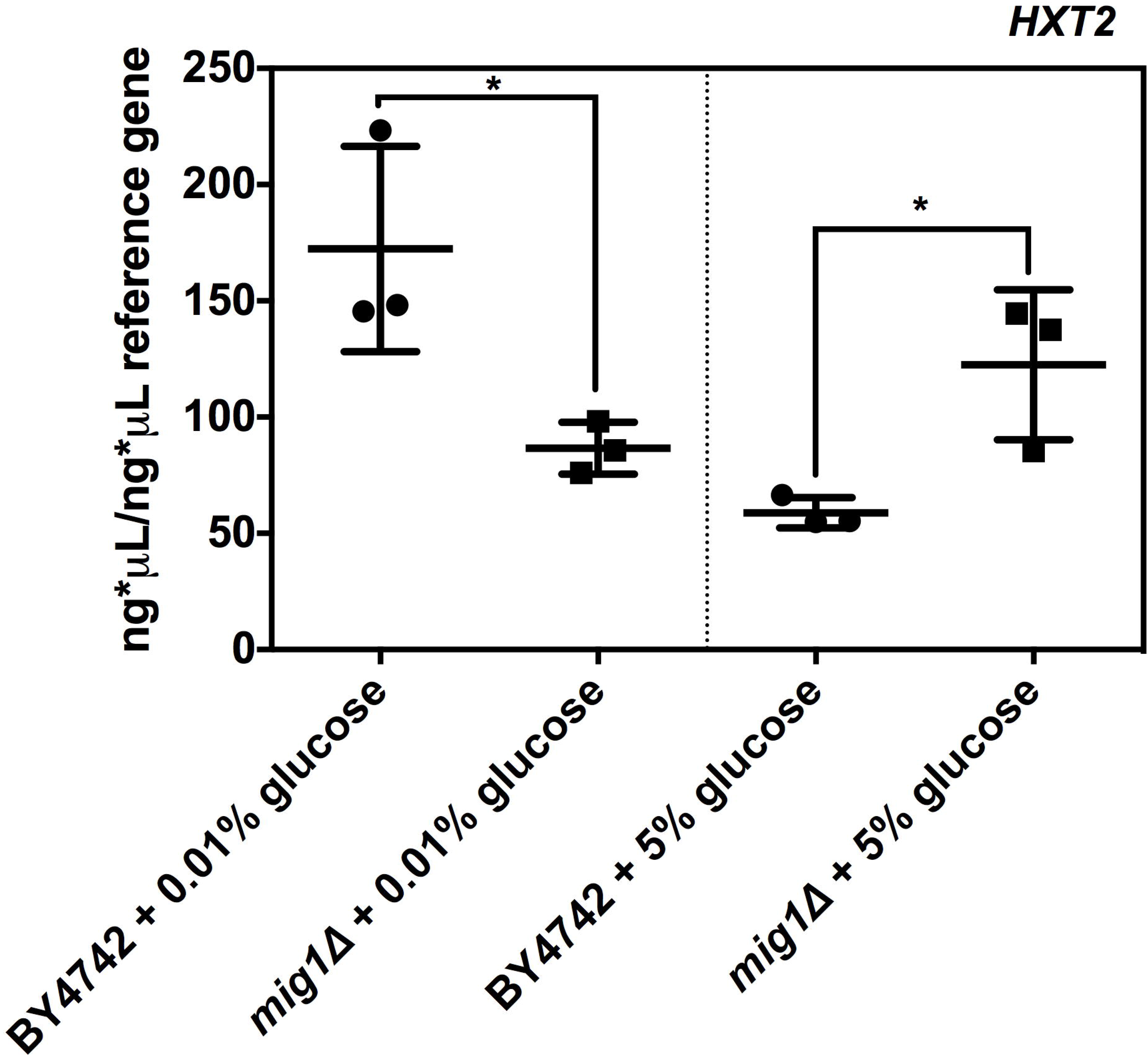
Influence of the *MIG1* deletion in the transcription of the *HXT2* gene at 0.01% and 5% glucose. Absolute quantification was made using a *UBC6* as a reference gene, from *S. cerevisiae* cultures in which pre-inoculum was grown with glucose in a YPD medium. The data represents mean ± standard deviation of three independent experiments with two technical replicates. Means were compared with a two-tailed unpaired *t*-test (\**P* < 0.05).

### Deletion of the genes *RAG5* and *SNF1* upon the growth of *K. marxianus*

The regulation of the pathway *SNF1/HXK2/MIG1* upon hexose transport transcription, exponential growth, and mitochondrial respiration suggests that this pathway could regulate the switch between mitochondrial respiration and fermentation; this arises the question: whether the pathway *SNF1/HXK2/MIG1* has a different role between Crabtree positives and Crabtree negative yeasts? To answer the last question, we used *K. marxianus*, which is a Crabtree negative yeast, that does not have the complex I of the electron transport chain, like *S. cerevisiae.* The orthologous genes of *SNF1* and *HXK2* of *S. cerevisiae* in *K. marxianus* are *SNF1* and *RAG5*, respectively. At 0.005% glucose deletion of the *SNF1* gene decreased the growth of *S. cerevisiae*, while in *K. marxianus SNF1* deletion increased the growth (**Fig. 13a**). In the case of the *HXK2* deletion did not have any effect on the growth of *S. cerevisiae* at 0.005% glucose, meanwhile, *RAG5* deletion augmented the growth of *K. marxianus* (**Fig. 13a**)*. SNF1* gene deletion lessened the growth of *S. cerevisiae* at 5% glucose, likewise, *Kmsnf1*Δ showed a minor growth than the parental strain of *K. marxianus* at 5% glucose (**Fig. 13b**). At 5% glucose *schxk2*Δ and *kmrag5*Δ strains displays a decrease in the growth of *S. cerevisiae* and *K. marxianus*, respectively (**Fig. 13b**). Overall, these data suggest the opposite role in the growth of the pathway *SNF1/RAG5* in *K. marxianus* promoting the growth at low-glucose concentration and decreasing the growth in high-glucose concentration. Additionally, at 0.005% is evident the divergent role on the growth of the pathway *SNF1/HXK2-RAG5* between *S. cerevisiae* and *K. marxianus.*

**Fig 13.**
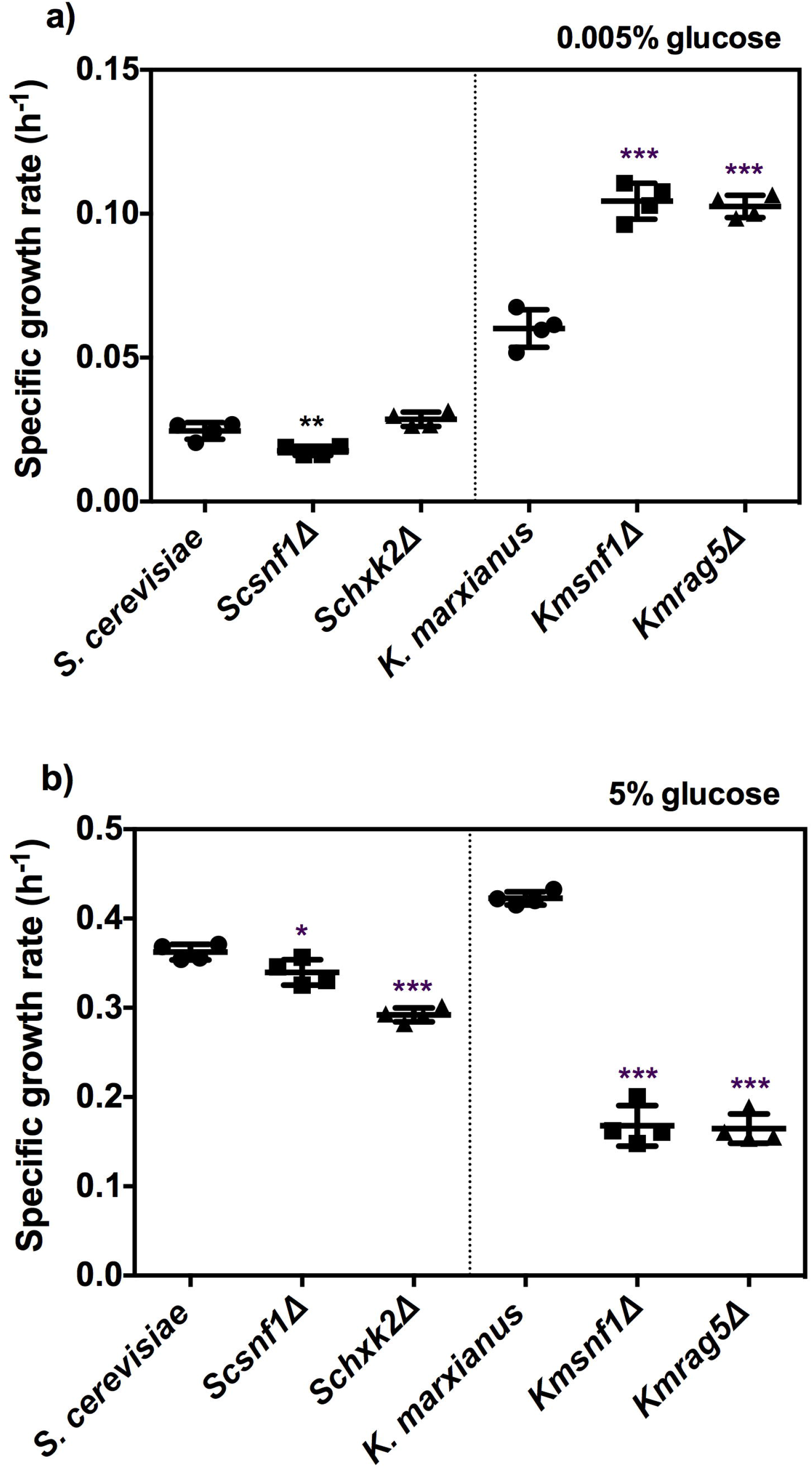
Influence of deletion of the genes *SNF1* and *HXK2/RAG5* in the growth of *S. cerevisiae* and *K. marxianus.* The exponential growth was represented trough the specific growth rate of the strain deleted in the genes *SNF1* and *HXK2.* In the case of *S. cerevisiae* the strains were named as *Scsnf1*Δ and *Schxk2*Δ, while for *K. marxianus* were designated as *Kmsnf1*Δ and *Kmrag5*Δ. a) Represents growth at 0.005% glucose; b) represents growth at 5% glucose. The data represents mean ± standard deviation of four independent experiments with two technical replicates. Means were compared using one-way ANOVA followed by a Dunnett multiple comparison *vs. S. cerevisiae* and *K. marxianus* (\**P* < 0.05; \*\**P* < 0.01;\*\*\**P* < 0.001).

### Impact of the via *SNF1/RAG5* pathway on the mitochondrial respiration of *K. marxianus*

To better understand how the pathway *SNF1/RAG5* impacts in the glucose-sensing of *K. marxianus*, that not presents the Crabtree effect, we measure its mitochondrial respiration. Interestingly, *K. marxianus* displays a diminution of mitochondrial respiration with the augment of the glucose concentration such as occur in *S. cerevisiae* (**Fig. 14a)**. However, the difference in basal mitochondrial respiration between *K. marxianus* and *S. cerevisiae* are 14.5 times higher at 0.005% glucose and 105 times higher at 5% glucose (**Fig. 14a)**. This result suggests that *K. marxianus*, although it is not a Crabtree positive yeast, responds similarly to glucose, lessening its mitochondrial respiration with lower levels of glucose.

**Fig 14.**
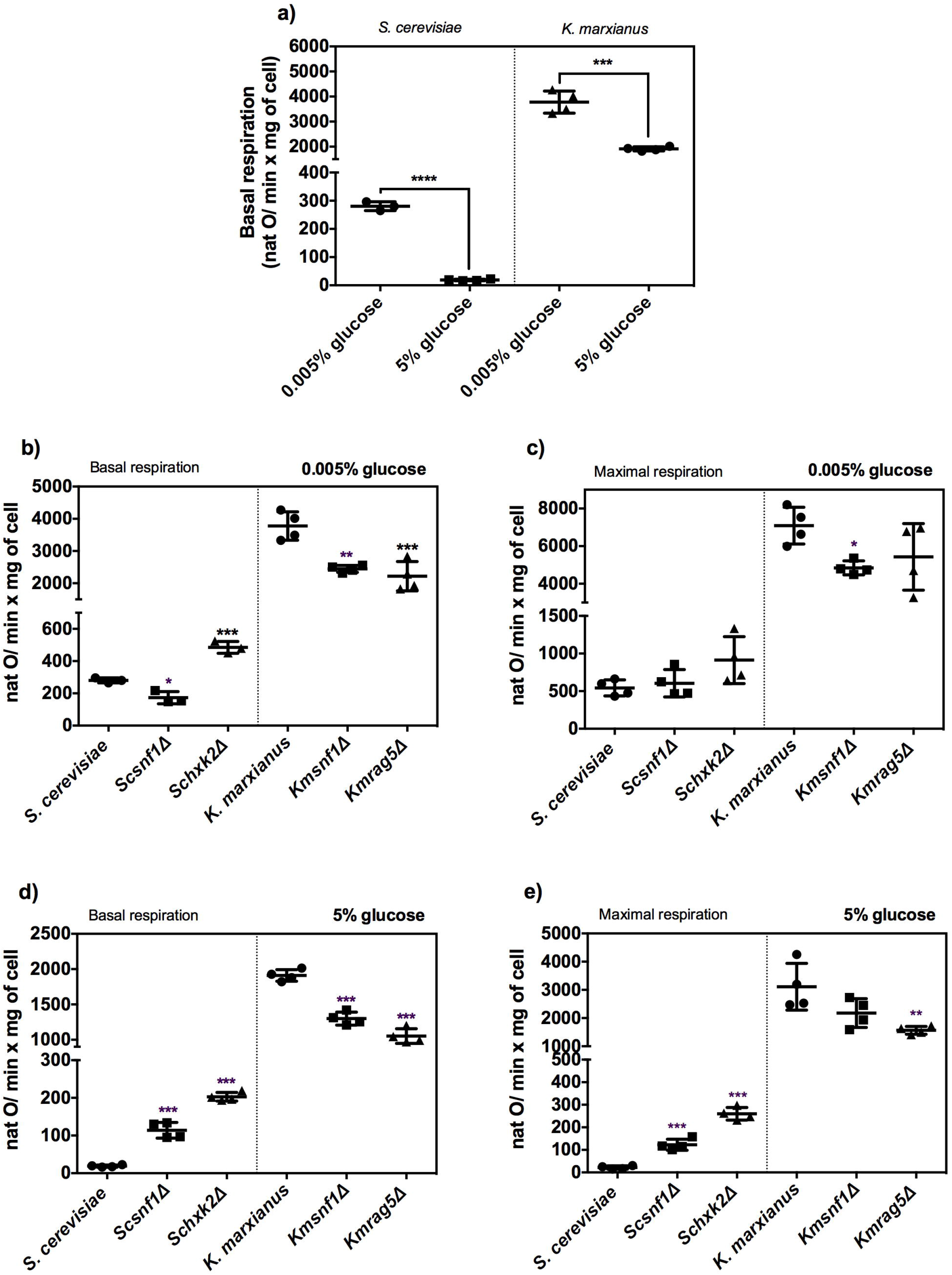
Effect of deletion of the genes *SNF1* and *HXK2/RAG5* in the mitochondrial of *S. cerevisiae* and *K. marxianus* at 0.005% and 5% glucose. Basal and maximal respiration were obtained from cultures in which pre-inoculum was grown with glucose in a YPD medium. a) Basal respiration of *S. cerevisiae* and *K. marxianus* strains at 0.005% and 5% glucose; b) basal respiration of the strains *S. cerevisiae* and *K. marxianus* and its deletant strains in the genes *SNF1* and *HXK2/RAG5* at 0.005% glucose; c) maximal respiration of the strains *S. cerevisiae* and *K. marxianus* and its deletant strains in the genes *SNF1* and *HXK2/RAG5* at 0.005% glucose; d) basal respiration of the strains *S. cerevisiae* and *K. marxianus* and its deletant strains in the genes *SNF1* and *HXK2/RAG5* at 5% glucose; e) maximal respiration of the strains *S. cerevisiae* and *K. marxianus* and its deletant strains in the genes *SNF1* and *HXK2/RAG5* at 5% glucose. The data represents the mean ± standard deviation of three independent experiments. For panels b), c), d), and e) means were compared using one-way ANOVA followed by a Dunnett multiple comparison *vs. S. cerevisiae* or *K. marxianus* (\**P* < 0.05; \*\**P* < 0.01; \*\*\**P* < 0.001). For panel a) means were compared with a two-tailed unpaired *t*-test (\*\*\**P* < 0.001; \*\*\**P* < 0.001).

The deletion of the *SNF1* gene has the same influence on mitochondrial respiration, decreasing it with respect to its parental strains at 0.005% glucose (**Fig. 14b-c**). In the case of the *HXK2/RAG5* deletion, there was an increment in mitochondrial respiration in *S. cerevisiae* and diminishing in *K. marxianus* when both were grown at 0.005% glucose (**Fig 14b-c**). At 5% glucose, the deletion of the *SNF1/HXK2-RAG5* genes has contrasting effects: increasing mitochondrial respiration in *S. cerevisiae* and decreasing mitochondrial respiration in *K. marxianus* (**Fig 14d-e**). These results highlight the opposite role of the *SNF1/HXK2-RAG5* pathway under high-glucose concentration that exerts the Crabtree effect, suggesting its participation in the molecular mechanism. Even, at 0.005% glucose *HXK2-RAG5* deletion also had a differential influence upon mitochondrial respiration.

## Discussion

The Crabtree effect mediated by glucose repression is a metabolic phenomenon with biomedical and biotechnological importance [27]. For example, in *S. cerevisiae* the ethanol production also occurs during the Crabtree effect under oxygen presence [28, 29]. The Warburg effect metabolic analogous to the Crabtree effect is present in the majority of the cancerogenic cells and its part of the cancer etiology [2, 30]. Nonetheless, the key piece to explain the molecular mechanism of the Crabtree effect is maintained elusive. Glucose transport could play an important role in the modulation of the glycolytic flux since it increasing is essential to induce the Crabtree effect. The wide-range of biochemical characteristics of the hexose transporters of *S. cerevisiae* implies a highly adapted metabolism to the uptake of sugars in different circumstances. However, the metabolic control associating hexose transporters expression with the Crabtree phenotype is not totally explained yet. The pathway Snf1p/Hxk2p/Mig1p has been associated with the transcriptional regulation of the hexose transporters [12, 16–18, 31]. Besides, the pathway Snf1p/Hxk2p/Mig1p has also been identified in the modulation of some phenotypes associated with the Crabtree effect, such as mitochondrial respiration [10], glycolytic flux [10], and fermentation [12]. Nevertheless, it has not been directly identified the direct genetic regulation of the hexose transporters with modulation of the Crabtree effect phenotypes by Snf1p/Hxk2p/Mig1p pathway. For this reason, the aim of this study was to identify the association between the transcriptional levels of hexose transporters with the growth and mitochondrial respiration, modulated by the Snf1p/Hxk2p/Mig1p pathway. In view of this, we provide evidence that the deletion of the *SNF1* and *HXK2* disturbs the exponential growth of *S. cerevisiae* in a glucose-dependent manner. Besides, *snf1*Δ and *hxk2*Δ strains impact in the mitochondrial respiration of *S. cerevisiae* in a glucose-dependent manner. The transcriptional levels of hexose transporters were also affected by the deletion of the *SNF1* and *HXK2* with a differential pattern in response to glucose concentration. A positive correlation was found between exponential growth and *Vmax* of the hexose transporters mainly transcribed; while a negative correlation was detected between mitochondrial respiration and *Vmax* of the hexose transporters mainly transcribed. Transcription of the gene *HXT2* was the most affected by the deletion of the pathway *SNF1/HXK2/MIG1* in the glucose concentrations tested (0.005%, 0.01%, and 5%). Finally, the deletion of the orthologous genes *SNF1* and *HXK2* in the Crabtree negative yeast, *K. marxianus,* has a differential effect in exponential growth and mitochondrial respiration in comparison with *S. cerevisiae* (Crabtree positive). In general, these results indicate that the *SNF1/HXK2/MIG1* pathway regulates the transcriptional levels of hexose transporters and this modulates the exponential growth and mitochondrial respiration.

Mitochondrial respiration and fermentation are pathways that enable microorganism division by ATP formation. It is expected that the alteration of the exponential growth rate, allow screening changes in bioenergetic metabolism [5]. Intriguingly, the increase in growth is a phenotype well documented during the Crabtree effect and with its analogous the Warburg effect, this higher growth occurs mainly under fermentation, which produces 2 mol of ATP per mol of glucose, 9 times lower than mitochondrial respiration in *S. cerevisiae* and 15-18 times lesser than mammalian cells. To maintain this growth and its concomitant ATP demands, the cells should be uptaken higher amounts of glucose through its hexose transporters. The glucose repression regulates those changes in the metabolism through the Hxk2p and Mig1p, repressing the transcription of genes that participates in the respiratory energetic metabolism [1]. In the case of Snf1p is a kinase that inhibits Hxk2p and Mig1p by phosphorylation, playing a role as a negative regulator of glucose repression. In the first part of this study, we evaluate the exponential growth, with the basis that changes in the energetic metabolism should impact in this type of growth. *SNF1* and *HXK2* gene deletions modified the exponential growth of *S. cerevisiae* with important differences. The *SNF1* gene deletion increases the exponential growth at 0.005% glucose (**Fig. 1b**) when it came from an inoculum grown with SC medium, whereas the same gene deletion decreased the exponential growth at 0.005% glucose when it came from an inoculum grown with the YPD medium (**Fig 13a**). A similar phenotype was observed with *HXK2* gene deletion, which increasing exponential growth when cells were precultivated in SC medium (**Fig. 1b**) and no change the exponential growth when the pre-inoculum was made with YPD medium (**Fig. 13a**). These data highlight the dependence of the exponential growth exerted by *SNF1* and *HXK2* genes deletion upon nutrient availability. The response of mitochondrial respiration to both *SNF1* and *HXK2* genes deletion was similar to the phenotype observed with exponential growth. At 0.005% glucose *snf1*Δ strain augments basal respiration when cells were precultivated in SC medium (**Fig 3b**) and the basal respiration was lessen in cultures coming from a pre-inoculum grown in YPD medium (**Fig. 14b**). The *hxk2*Δ strain did not change the basal respiration at 0.005% glucose (**Fig 3b**) when it came from a pre-inoculum grown with SC medium, while increases the basal respiration at 0.005% glucose when cells were precultivated with YPD medium (**Fig. 14b**). These results remark the association of the exponential growth with mitochondrial respiration and also underline the intricate involvement of *SNF1/HXK2* with the nutrient disposition in the media, highlighting the importance of describing the pre-inoculum to the better comparison of data.

As expected, the *HXK2* gene deletion reverts two important phenotypes observed during the Crabtree, diminishing the exponential growth (**Fig. 2d and 13b**) and increasing the mitochondrial respiration both at 5% glucose (**Fig. 3d and 14d**). A diminution in the exponential growth through the doubling time has also been reported, in the *hxk2*Δ strain grown in YPD medium and SD minimal medium coming from *S. cerevisiae* DBY2230 genetic background [32]. Since *SNF1* has opposite participation in the glucose repression to *HXK2*, we expected to observe the phenotypes related to the Crabtree effect unchanged. The *snf1*Δ strain precultivated in the SC medium did not change either exponential growth (**Table 3**) and mitochondrial respiration (**Fig. 3d**). However, *snf1*Δ strain pre-inoculated in the YPD medium showed a slight decrease in exponential growth (**Fig. 13b**) and an increase in mitochondrial respiration at 5% glucose (**Fig. 14d**). The increase in the mitochondrial respiration was also observed in *snf1*Δ cells grown in SC medium with 2% glucose [12] and in *snf1*Δ cells grown in YPD medium supplemented with 10% glucose [10]. These phenotypes make clear the participation of the Snf1p at high-glucose concentrations, since it impairs the mitochondrial respiration. Interestingly, Snf1p regulates the arresting◻ related trafficking adaptor Rod1p/Art4p, which mediates the endocytosis of the high◻affinity glucose transporter Hxt6p at high glucose concentrations [31]. Besides, the *SNF1* gene lacking stimulates Hxt1p and Hxt3p endocytosis and degradation in the vacuole [33]. Importantly, overexpression of low-affinity transporters Hxt1p and Hxt3p rescues *snf1*Δ growth supplemented with 0.01% 2-deoxyglucose [33]. These studies remark the strength relation between *SNF1* and the hexose transporters expression. Additionally, these results indicate the participation of the *SNF1/HXK2* in the exponential growth and the mitochondrial respiration, two important phenotypes observed in the Crabtree effect.

*S. cerevisiae* count with a numerous family of hexose transporters that have a wide range of biochemical characteristics and is according to its affinity (*km*) that hexose transporters were classified [34]. However, little attention has been putting in the *Vmax*, which describes the capacity of each hexose transporter to uptake sugar in a unit of time. Remarkably, a positive correlation has been found between the sugar uptake rate and ethanol production rate in *S. cerevisiae* [7, 8], which suggests that the key piece in the establishment of the Crabtree effect might be the glucose transport. Here, we found that exponential growth (**Fig. 7a and Fig. 10**) and mitochondrial respiration (**Fig. 7b**) correlates with the *Vmax* of the mainly transcribed hexose transporters. These results reveal that rate of glucose uptake (*Vmax*) could be an important mediator in the establishment of the phenotypes (augmented growth and mitochondrial respiration repressed) related to the Crabtree effect. In support of this idea, a linear correlation between the natural logarithms of the *Vmax* of hexose transporters and the glucose consumption rate in *S. cerevisiae* was found [7]. In addition, when *S. cerevisiae* cultures grown in aerobic chemostat using ethanol as a carbon source were exposed to glucose (fructose) pulse, the strain that only expressed Hxt1p transporter changed its phenotype to a respiro-fermentative metabolism as did the wild-type strain, whereas strain expressing Hxt7p transporter maintained respiratory metabolism [35]. Importantly, the PCA revealed that *HXT2* transcription was the most affected by the *SNF1/HXK2/MIG1* (**Fig. 11**), which might have important effects on the phenotypes exerted by this pathway. Overall, these results highlight the relationship between hexose transporters activity, glucose uptake rate, and mitochondrial respiration/fermentation. Additionally, our results reveal that the *SNF1/HXK2/MIG1* pathway regulates in a transcriptional way the hexose transporters, this pathway also showed a linear correlation with the exponential growth and mitochondrial respiration, two characteristic phenotypes presented during the Crabtree effect.

## Funding and additional information

This work was supported by the grant: PROMEP program (ITESCH-EXB-002) awarded to LAMP that includes the scholarship grant to JGMG.

## Conflict of interest statement

The authors declare that they have no conflicts of interest with the contents of this article.

## Notes

### Competing Interest Statement

The authors have declared no competing interest.

